# Transferable Collective Variable to accelerate Protein-Ligand (Un)Binding Transitions via Explainable Machine Learning and Intriguing Role of Ligand Solvation

**DOI:** 10.64898/2026.08.21.746233

**Authors:** Saikat Dhibar, Biman Jana

## Abstract

The process of drug unbinding is of immense importance in the field of biophysics and therapeutics. The behavior of these systems is greatly influenced by their thermodynamic and kinetic properties. Therefore, it is crucial to accurately estimate the ligand binding free energies and rate of ligand dissociation, yet these processes are often governed by rare event transitions that lie beyond the reach of standard brute-force molecular dynamics simulations. While enhanced sampling simulations offer a solution, their efficacy is strictly contingent upon the selection of appropriate collective variables (CVs) which is non-trivial for complex systems like protein-ligand complexes. In this study, we present a method to derive optimized CV from transition state region (TS) via an interpretable machine learning (ML) model, Elastic Net. By employing some physically intuitive order parameters, the derived optimized CV from the TS-region greatly accelerate ligand binding-unbinding transitions and achieves rapid free energy surface (FES) convergence across diverse systems including buried and solvent exposed active sites such as Trpsin-benzamidine complex, host-guest systems and sodium epoxidase etc. Intriguingly significant contribution of the ligand hydration is found in the optimized CV which depicts crucial role of solvent in driving ligand binding-unbinding transitions. The estimated binding free energies for different protein-ligand complexes match quite well with experiments, while maintaining a low computational cost. The derived optimized CV is also used to calculate the ligand residence times across different systems and calculated residence times are within the experimental range for all systems, again with very little computational costs. Moreover, we show that the optimized CV constructed from TS region via an interpretable ML model is transferable across diverse systems, offering a robust and scalable framework for drug discovery and investigation of complex biomolecular recognition.

## Introduction

The molecular recognition between a protein and its ligand is a fundamental process governing the majority of biological functions and pharmacological responses.(1) In the realm of modern computer-aided drug design, the ability to accurately calculate thermodynamic parameters, such as binding free energy and kinetic properties, such as the ligand dissociation rate is extremely important.(2, 3)Atomistic molecular dynamics simulations provide a high-resolution microscopic lens into these processes which shed lights on the mechanism of binding-unbinding transitions and allosteric pathways in biomolecules.(4–8) These simulations also used routinely to estimate absolute binding free energies and rates associates with these processes. However, the accuracy of these estimates largely governs by the quality of the sampling. The multiple transitions between different metastable basins are required to achieve good estimates of these parameters. This present a sampling bottleneck for ligand binding-unbinding transitions as these transitions are rare events where the different metastable conformations are typically separated by high free-energy barriers that make spontaneous transitions extremely difficult to observe within standard brute-force simulation. To bypass these limitations, several enhanced sampling simulation methods have been developed over the last decades. Examples of these methods include umbrella sampling(9), Metadynamics (MetaD),(10) Adaptive biasing force(11), alchemical free energy calculations(12), thermodynamic integration(13, 14) and Gaussian accelerated molecular dynamics (GAMD)(15, 16). Besides these methods, adaptive sampling and MSM are also used to investigate drug dissociation process and calculate the kinetics associated with these processes.(4, 6, 17) In this study, we will frame our discussion in the context of MetaD and its variants as these methods are the major focus of this current work. In these methods an external bias potential along some pre-chosen collective variables (CVs) is applied which enforce the system to overcome large free energy barriers present between the metastable states.

Enhanced sampling methods are widely used to study several different biomolecular recognition processes like Protein Folding-Unfolding, ligand binding, coupled folding upon binding and many different biomolecular processes.(18–22) Deriving mechanistic insights from enhanced sampling simulations hinges on getting a converged free energy landscape for protein-ligand binding and unbinding. However, the success of this class of methods heavily relies on the carefully chosen CVs which should capture crucial slowest degree of freedom for ligand recognition processes.(23–25) Deriving such optimal CVs for complex bio-molecular processes like protein-ligand binding is a formidable task and requires extensive expert knowledge about the system of interest. Choosing a wrong CVs can significantly affects the sampling efficiency of the enhanced sampling methods, thereby limiting our understanding about the processes. Traditional intuitive descriptors, such as the simple distance between a ligand and its binding pocket, frequently fail to yield converged free energy estimates because they neglect critical slow processes such as hydration of the binding site, hydration of ligand, ligand orientations etc.(21, 26, 27)

In recent years, machine learning (ML) models emerged as a powerful tool to obtain optimal CVs for different bio-molecular process with no exception in ligand binding-unbinding transition. Over the last decades, significant advancements have been made particularly in the class of MetaD methods to accelerate the sampling of ligand binding transitions. Earlier, Limongelli et al. developed Funnel Metadynamics (FM) which imposes a funnel shape restrain potential on the ligand, addressing the entropic bottleneck present in sampling the unbound state.(28) Parrinello and co-workers developed VAC-MetaD to accurately estimating binding free energies and kinetics for protein-ligand complexes using variational principal where Trypsin-Benzamidine complex is used as a prototypical example.(29) Tiwari and colleagues proposed Reweighted Autoencoded Variational Byes (RAVE)(30)and applied this method to investigate ligand binding mechanism in hydrophobic cavity-fullerene system and L99A T4 lysozyme benzene complex.(30, 31)Expanding on this, they introduced State Predictive Information Bottleneck (SPIB) which utilizes an iterative procedure to learn and improve the CV via variational autoencoder (VAE) framework from set of input descriptors.(32)The integration of SPIB with infrequent metadynamics (iMetaD) has been used to estimate the drug dissociation kinetics of many protein-ligand complexes that align well with the experiments.(33) Rizzi et al. identified different hydration spots in the binding site and combine them to derive an optimized CV through Linear Discriminant Analysis (LDA) to estimate binding free energies in different host-guest systems from SAMPLE 5 blind challenge of binding free energy prediction.(21) Gervasio and co-workers have estimated absolute binding free energies for multiple protein-ligand complexes of heat shock protein 90 (HSP90) and bromodomain-containing protein 4 (BRD4) with their own developed method, One On-The-Fly Probability Enhanced Sampling (OneOPES) which uses multiple geometric CVs simultaneously with different methods of OPES and replica exchange ladder.(34)Very recently, Elangon et al. have calculated the absolute binding free energies in different host guest systems via combined protocol of OPES-MetaD and OPES-EXPLORE which agree well with reference values.(35) They have also developed a method which can identify mechanistic pathways of ligand binding using multi-task learning framework via well tempered metadynamics (WT-MetaD) and artificial neural network (ANN).(36) Besides these methods, there are plenty of studies which provides mechanistic insights into the ligand binding phenomena via different advanced simulation protocols.(37, 38)

These methods significantly address the problem of selecting appropriate CVs to investigate ligand binding phenomena and enhance our understanding about the recognition processes. While effective, these methods often ignore the crucial hydration CVs of ligand or binding site. Moreover, the cost of deriving optimized CV and running enhanced sampling with them is sometimes high, particularly for ML-derived CVs. In addition to that, these methods are designed to derive optimized CVs each time for a new system which broadly limits their applicability in real drug discovery pipeline. In this manuscript, we propose a method based on explainable artificial intelligence (XAI) model to derive optimized CVs from transition state ensemble (TSE) via choosing some set of physically intuitive descriptors to investigate range of protein-drug complexes. The derived CV from TSE rapidly accelerates the ligand binding transitions in both solvent exposed and buried active site complexes. The computed absolute binding free energies from this method match notably well with the experimental values. Moreover, the method is integrated with infrequent metadynamics (iMetaD) to calculate drug residence times for each system studied here and encouragingly residence times are also within reasonable ranges with experiments. The interpretable nature of the method identifies ligand solvation as the major driving force in recognition phenomena compared with that of active site solvation. Finally, the transferability of this method is checked which is often overlooked and a transferable CV is proposed to investigate the ligand binding mechanism in any complexes. This overcomes the need of new training data for the development of any ML-CVs in new protein-ligand complexes and bypasses the knowledge of system specific input descriptors. Overall, by merging TS information with explainable AI model, the transferable CVs proposed here provides a robust approach for navigating the complex biomolecular landscapes essential to modern drug discovery.

## 2. Methods

In this section we gave a short description about different method used in this work. This includes well tempered metadynamics, infrequent metadynamics, Time lagged denoising autoencoder, Elastic Net and details of running MD simulations. The details iMetaD and funnel metadynamics setup are given in SI note SI.

### (A) Well-tempered metadynamics (WT-Metad) and infrequent MetaD

Metadynamics (Metad) is a one of most popular enhanced sampling method for investigating rare event processes.(10, 39) Metad enhances sampling efficiency by applying a history-dependent bias which is constructed as a gaussian function to some specific degrees of freedom in the system which is referred as CVs. These CVs should incorporate relevant slow degree of freedom of the systems. In well-tempered metadynamics the height of the Gaussian gradually decreases with time which allows faster free energy convergence. The MetaD potential helps to system to escape from the stable minima, overcome free energy barriers and explore FES within reasonable computational cost. Moreover, in MetaD after a short time span the simulation enters a quasi static regime in which expectation value of any observable can be estimated by a running average.(40) From the WT-MetaD, FES as a function of some CV which is directly biased in MetaD or some other coordinate which was not directly used in biasing purpose can be obtained from a reweighting procedure.

In infrequent MetaD, the rate at which bias is added is reduced in such a way that the possibility of adding bias in TS becomes low.(41) Quite intuitively, if the rate of bias deposition becomes extremely low, the simulation would tend to approach unbiased MD with the cost of extremely slow acceleration of sampling. With appropriate approximations, unbiased timescale of the processes can be estimated from this infrequent MetaD run by multiplying the biased timescale with accurate acceleration factor.(42) The reliability of the timescales so obtained can be quantitatively assessed through a Kolmogorov–Smirnov (KS) test.(43) The details mathematical equations to estimate kinetics from iMetaD runs are given in SI note S1.

### (B) Time-lagged denoising autoencoder

This framework utilizes adenoising autoencoder (DAE) architecture tolearn the relationship between input and output via compressed bottleneck dimensions.(44)Unlike traditional autoencoder, DAEintroduces random noise to the input feature descriptors forcing the network to learn robust features by reconstructing the original, noise-free input. This framework tries to minimize overfitting which is a very common challenge in the development of any ML models. In our framework, a very small amount of noise is added into the input descriptors calculated from short iMetaD runs. The temporal information is integrated by incorporating the lag-time into this framework, drawing inspiration from methodologies like VAC-Metad(45), Deep-TICA(46) and VAC-encoder(47) approaches. This integration allows the network to learn the slowest motion associated with the process of interest. The architecture uses a symmetric autoencoder with three hidden layers in both encoder and decoder dimensions. For the encoder layer linear activation function is used for better interpretability and non-linear activation function, tanh, is used for the decoder layers.

Form the pool of iMetaD trajectories, a set of input descriptors are calculated and fed into the NN architecture in the framework of denoising autoenoder. The training procedure is designed to minimize a composite loss function which comprises of reconstruction loss as well as autocorrelation loss inspired by Ref. The mathematical forms of the loss functions are given in equation (1),(2) and (3).

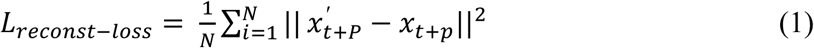

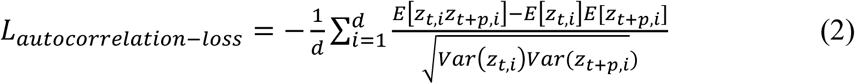

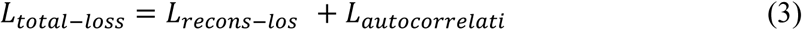

x_t_ is the input at time t. x_t+p_ is the time-lagged input after a lag time p. x_t+p_^’^ is the predicted time-lagged input from the decoder. Equation 1 describes the mean-squared-error (MSE) loss for clean time-lagged input descriptors constructed from noisy time-lagged input descriptors. In equation 2, d is the dimension of latent space, E and Var represents the sample mean and variance. Z_t,i_ is the i^th^component of latent dimension at time t and Z_t+p,i_ is the i^th^ component of latent dimension after a lagtimet+p. Equation 3 represents the total loss function of time-lagged denoising autoencoder model where both the loss functions are designed to find out slowest components from MD simulation data.

### (C) Elastic-Net surrogate model to identify the input feature descriptors importance

In recent times, explainable AI model has gained much attention and has been extensively used in wide range of biological problems.(48–52) In enhanced sampling community, explainable AI model is used to identify importance of OPs in the true reaction coordinate and also directly used to enhance the sampling of bio-molecular processes.(53–57) Previously, our group has identified the role of solvent in polymer collapse transition process via interpretable AI model, SHAP.(58) And also we have constructed optimized CV for polymer collapse transition processes via Elastic Net model to enhance the collapse transition in different length of polymer systems within a low computational cost.(22) In this work, we have used this protocol to more relevant and challenging problem of protein-ligand dissociation. We have generated multiple independent reactive trajectories going from bound to unbound conformations biasing a trial CV, COM distance between binding site and ligand. Here, we have moved on to EN model to assess the coefficients of feature descriptors locally. EN model is a combinations of both Lasso and ridge regression model with both l1 and l2 regularizations. Very recently Chatterjee et al. have demonstrated a surrogate model CV based on lasso regression that efficiently construct the underlying free energy landscape at low computational cost compared with the neural network.(57)Using EN model, we locally derive the coefficients of different OPs specifically at the TSE to accelerate the ligand binding-unbinding transition. In contrast, in the approach of Chatterjee et al.(57) The LASSO CV is constructed using latent variable derived from the time-lagged autoencoder model. In this study, the relation between input descriptors and output of EN model is described in the following form:

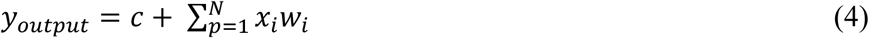

Here y_output_ denotes the predicted value from the model. c is the bias term and N is the total number of input feature descriptors,x_i_ is the value of i^th^feature descriptor and w_i_ is the coefficient of i^th^descriptor. In this work y_output_ is the committor probability values (ranging from 0.35 to 0.65) predicted from the model using three input OPs: funnel axis distance, ligand solvation and active site solvation. The bias and weights are optimized by the following loss function:

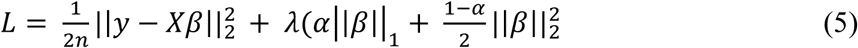

The first term in this equation 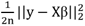 implies Mean squared error term. ***λ*** controls the regularization strength. α is the mixing parameter which controls how much weight is given to l_1_and l_2_regularization methods. Generally, it varies between 0 to 1 in the EN model. I|β|I is the l_1_ norm (Lasso) which provides sparsity and is the squaredl_2_norm (Ridge) which shrinks the coefficients but does not force them to exactly zero 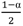 term ensures proper weighting and normalization with MSE loss.

### (D) Setup of MD simulation details

The host guest systems considered in this work, input parameter files are taken from the previous works.(21, 59) The simulation parameters and starting conformation are identical to that of the previous work. In a nutshell, systems are parameterized using GAFF force field using RESP derived charges and water molecules are modeled using TIP3P water model. The simulation is performed at 300 K via V-rescale thermostat with 2 fs timestep. We have taken the input parameter files for Trypsin-Benzamidine complex from reference 18.(26) The protein is modeled using Amber-14SB force-field and ligand interactions are taken from Amber GAFF library. TIP3P is used as a water model. We performed the simulation at 300 K temperature in the NPT ensemble with timestep of 2 fs. In case of hydrophobic cavity fullerene system, input files are taken from the github repository of Ref. 35.(54) System is solvated in TIP4P water model. Lennard-Jones interactions between particles are evaluated using Lorentz-Berthelot mixing rules and taken as it is from the Ref 35. For HSP90 ligand complex, input parameter files are taken from ref. 27.(34) Amberff14SB is chosen to model HSP90 protein.TIP3P water model is used to solvate the protein. Final simulation is performed in NVT ensemble in 300 K temperature. For soluable sodium epoxide hydrolase ligand complex and L99A T4 lysozyme benzene inputs are taken Ref. 47 and 48 respectively.(60, 61)For, sodium epoxide hydrolase ligand is parameterized using AMBER force field (GAFF2) with RESP charges and immersed in a box containing TIP4P-D water model. a99SB-disp force-field is chosen. For lysozyme complex interaction is modeled using CHARM22* force-filed and TIP3P water model is used. For both the cases final MD run is performed in NPT ensemble with 2fs timestep at 300 K temperature. More details about the simulation procedure are found in SI note S2.

All the MD simulations are performed for all the complexes using GROMACS 2020.6.(62)For running well-tempered metadynamics (WT-Metad) simulation, GROMACS 2020.6 is patched with PLUMED 2.7.4.(63) Sigma for WT-Metad is determined by calculating the standard deviation of the particular CV from their unbiased run. The chosen sigma for each system are given in Table S1.The height of the gaussian is kept fixed for a particular system irrespective of different CVs used for biasing for a fair comparison. The explicit details of biasfactor for each systems are given in Table S2.The bias is added with a deposition rate of 1/500 steps for each system. After performing WT-Metad simulations, we have reweighted the FES using Tiwari and Parrinello’s method.(64) The convergence of WT-Metad is checked by comparing the FES at different time intervals during the simulations.

### (E) Protocol

In this section we have described the methodology of constructing TS-derived CV in detail. For construction of any ML-CV, one needs training data generated from either biased or unbiased MD runs. Unfortunately, generating training data from long unbiased MD simulation is troublesome specifically for complex systems like protein-ligand binding. One needs to have multiple ultra long trajectories to observe one binding or unbinding events. To mitigate this issue, we have generated multiple transition paths between bound and unbound conformations using MetaD which are typically just 5-20 ns long. This approach drastically reduces the computational cost of have long trajectory data. Examples of generating training data for the construction of ML-CV from biased runs also reported in the previous literature. Moreover, to ensure the sampled TS in our initial exploratory runs is biasfree, we intentionally use infrequent MetaD where slow bias deposition rate decrease the possibility of adding bias in the TS region. As the optimal CVs for any new protein-drug complexes is not known beforehand, we run initial iMetaD with distance as an OP, in principal one can generate these initial trajectories with some other trial, suboptimal coordinate to sample the transition pathways between bound and unbound conformations. From these short iMetaD runs (typically ∼ 4-5), we have generated some reactive events going from bound to unbound state. Using these trajectories, some input descriptors are calculated and passed through a time-lagged DAE model which comprises of multiple hidden layers. From the time-lagged DAE, model compressed latent dimensions areobtained which capture dynamics of the systems. Then, the TS region is identified using the committor. A dataset comprises of normalized input descriptors along the committor values is constructed. In this dataset, normalized input descriptors serve as features and committor probability values serve as target variable. We have selected the transition state data points where committor probabilities are in the range of 0.35-0.65. Next, we have trained a linear ML model, EN with the normalized input descriptors to predict committor values in the TS region. 5-fold cross validation is used to find out the optimum hyperparameters values of EN model. A sensitivity analysis was conducted in which the committor windows were shifted, allowing us to evaluate the dependence of the EN coefficients on the selected committor ranges and results is shown in igure S1.With these tuned hyperparameters of EN model, we have calculated the coefficients of OPs near the TS region. In the end, we have constructed the TS-derived-CV by linearly combining the weights of the OPs and run final MetaD runs with this TS-derived CV to explore the FE . The workflow of the proposed method is shown in Figure 1.

**Figure 1:**
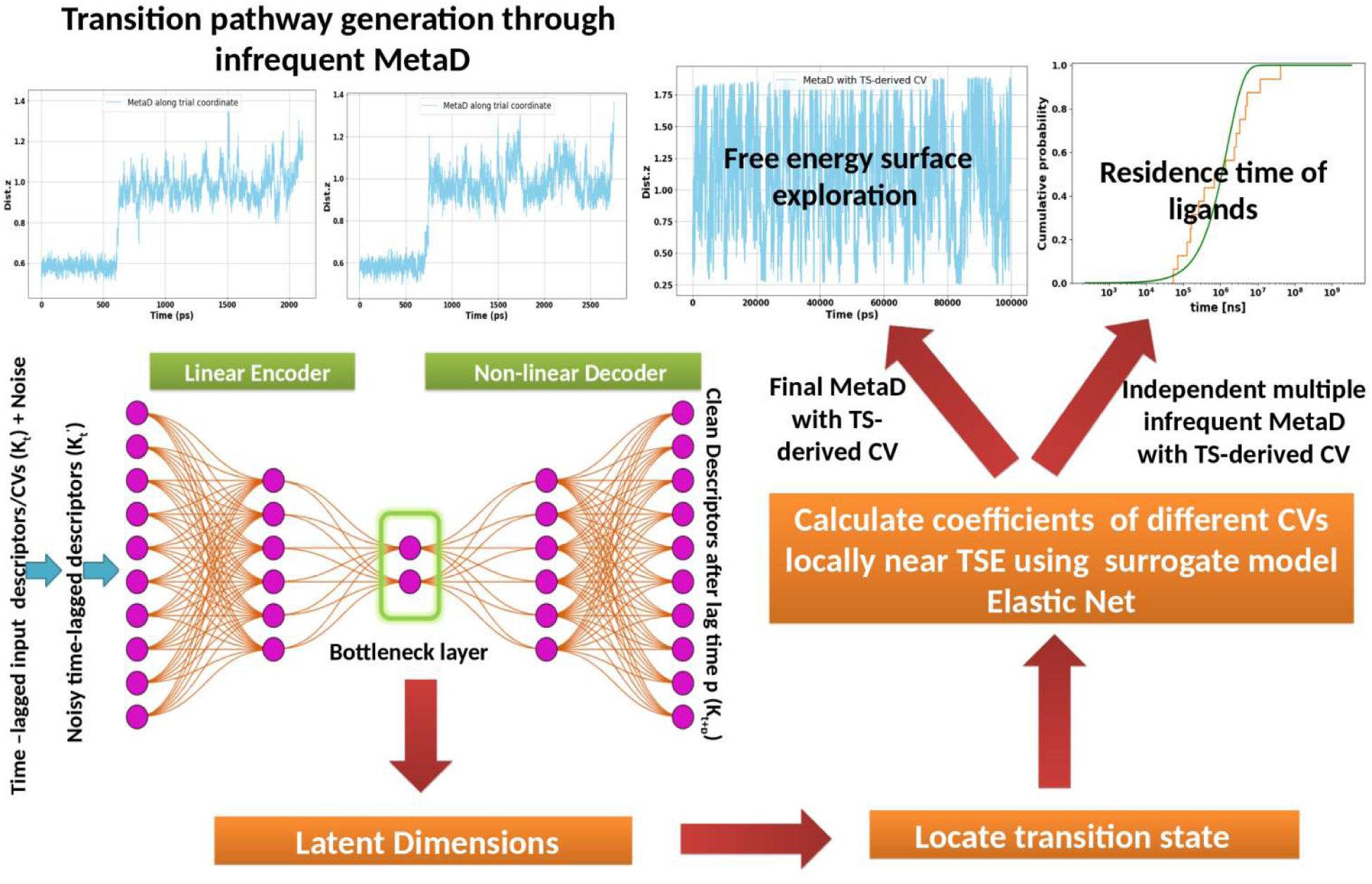
Overall workflow of the proposed protocol for running MetaD to investigate ligand binding/unbinding transitions. Starting from the bound conformation, a few reactive trajectories are generated via a trial OP. Then some descriptors are calculated from these trajectories and feeding them through a DAE model to get the compressed latent dimensions. From these latent dimensions, TS is located and relevance of different input feature descriptors are estimated via interpretable ML model. Finally a new CV is constructed with the estimated weights to run final MetaD to explore FES and estimate kinetics with independent iMetaD runs.

## 3. Results and Discussion

### A. Case of solvent-exposed binding-site

We have selected five different systems which has solvent exposed binding site. To construct the TS-derived CV for solvent-exposed case we have selected three OPs namely: (a) funnel axis distance, (B) active site solvation and (C) ligand solvation. Then we have evaluated the coefficients of these OPs near TS region using EN model as described in the methods section. The normalized coefficients derived from EN model for Trypsin-Benzamidine complex is tabulated in Figure 2(A). Strikingly, we find that ligand solvation plays the most dominant role in final TS-derived CV, followed by active site solvation. The role of active site hydration is investigated by other groups in the past, however the striking role of ligand solvation is not well described in the past. Funnel axis distance has the lowest contribution in the optimized CV, but not negligible. This highlights distance based geometric OPs has relatively minor influence in facilitating ligand binding transition. We have evaluated relevance of these three OPs near TSE for each system. The normalized coefficients near the TSE for the other three systems are shown in Figure S2. Quite interestingly we have found that ligand solvation emerges as the most dominant coordinate for each case, followed by active site solvation and distance CV. While the coefficients of these OPs near TSE vary a little, relative order remains consistent. In Figure 2B, we have given the bound conformations of systems studied in this study to investigate ligand binding in solvent exposed cases.

**Figure 2:**
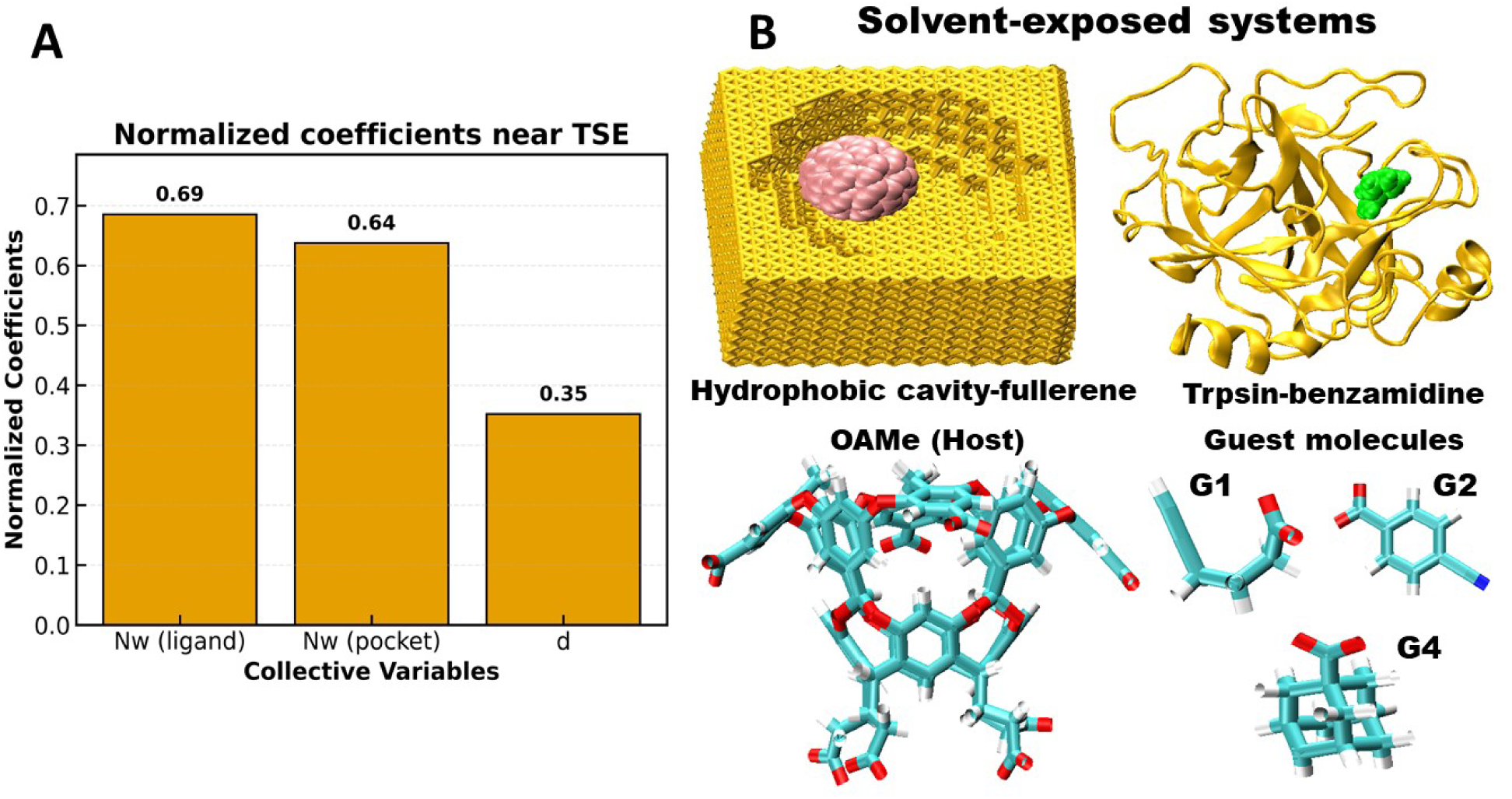
(A) Normalized coefficients of different OPs near TSE for Trypsin-Benzamidine system. (B) Bound conformations of protein-ligand complexes used in this work for solvent exposed active site. Structures are shown using Visual molecular dynamics (VMD)(65) using New Cartoon, VDW and Bonds representations.

As it is already mentioned, CVs importance near the TSE remains consistent across the different solvent exposed systems. It highlights crucial role of both ligand and active site solvation in driving the ligand binding transitions. Therefore, it prompted us to test the transferability of the constructed CV. We have used TS-derived CV for Trypsin-Benzamidine to directly apply to investigate the ligand binding mechanism in other solvent exposed systems. Subsequently, we run final MetaD straight way with the TS-derived CV where coefficients are obtained from Trypsin-Benzamidine complex system. Next, we check the quality of the constructed TS-derived CV in FES exploration as well as free energy convergence. One of key quality of a good CV in enhanced sampling simulation is that the CV should trigger multiple back and forth transitions between the metastable basins.

Hydrophobic cavity-fullerene system is one of the model system to study ligand binding mechanism and many previous studies made effort to understand the binding mechanism in the hydrophobic cavity previously.(54, 66, 67) We have shown the time series plot of distance over 200 ns in Figure S2 where almost 15 binding-unbinding transitions are observed. The reweighted FES along distance and ligand solvation is shown in Figure S3. It is shown there is a large barrier of ∼ 80 KJ/mol present going from bound and unbound conformations consistent with the previous studies. The FE convergence plot shown in Figure S4 depicts after just 66 ns FE reaches a plateau.

Next we have tested our methodology in host-guest complexes of SAMPLE5(68, 69) challenge which are used by other enhanced sampling methods to test the efficacy in past.(21, 59, 70) This system consists of a octa acid calixarene host (OAMe) with different ligands. We have investigated the ligand binding mechanism of OAMe host with three different ligands namely: G1, G2 and G4. The results for G1 ligand are shown in Figure 3A. The time series plot of dist (z) of the WT-MetaD run of OAMe-G1 shows large number of transitions between bound and unbound conformation. The reweighted FES shown in Figure 3A, depicts that there are three metastable basins present (corresponding configuration are also shown in the figure): bound and unbound and bound-like intermediate state, where in intermediate state G1 ligand is slightly upwards compared with that of original bound conformation. The main FE barrier of ∼16 KJ/mol is present going from intermediate state unbound conformation. We have also run WT-MetaD with funnel axis distance and presented the results in Figure S5(A-B). It depicts in the free energy convergence plot that the FE is converged after 150 ns. However, biasing with only funnel axis distance does sample the bound like intermediate properly as compared with TS-derived CV. The results for OAME-G2 are presented in Figure S6. The FE convergence plot shown in Figure S6B depicts that FE converges within 70 ns. The results of OAMe-G2 biasing with distance CV is given in Figure S7, where it took a slightly longer time to converge. In Figure 3B, the results (time series, FES, and configurations of different states) for OAMe-G4 are presented. Interestingly, for OAMe-G4 complex, reweighted FES in Figure 3B shows intermediate state is almost equally stable with that of bound conformation. Due to the presence of this crucial intermediate state, FES exploration for this complex is much more difficult than the other host-guest systems. The major barrier for this complex is present between the bound and intermediate state of about 13 KJ/mol. Previous studies have converged the FES for this system using a greater number of OPs and employing more sophisticated enhanced sampling method coupled with multiple replicas. WT-MetaD biasing with only funnel axis distance is performed and free results are presented in Figure S7 where it took longer time to converge the FE than TS-derived CV. It is important to note that, a huge number of re-crossing events between the bound and unbound conformations are observed for each system within just 200 ns using our TS-derived CV as a biasing variable. FE convergence plot for both the complex shown in Figure 3(D-E) depicts that FE along dist.z reaches a plateau after ∼ 60 and 72 ns respectively. The required FE convergence times using TS-derived CV is quite lower compared to the previous studies.

**Figure 3:**
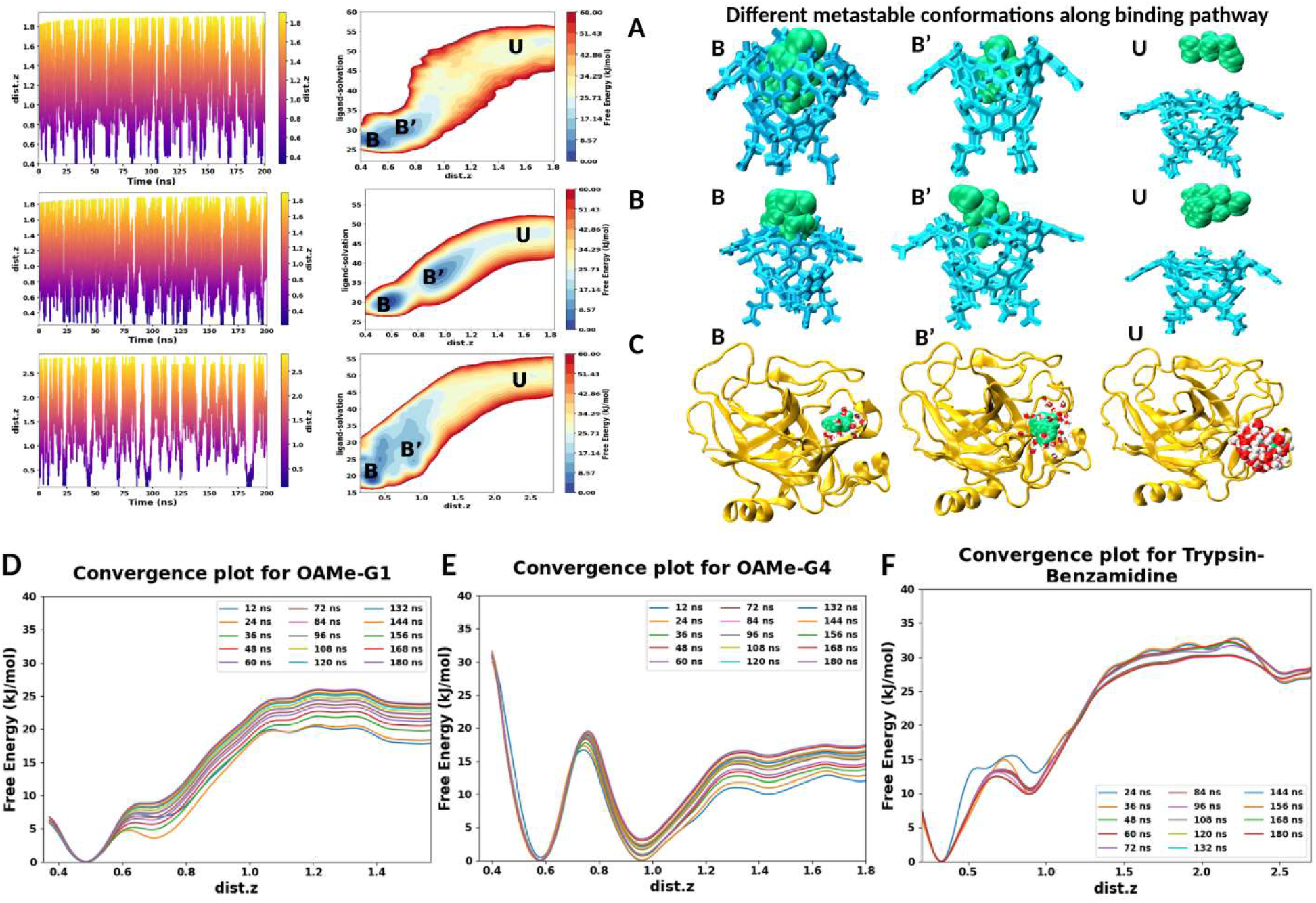
(A-C) Trajectories of WT-MetaD biased with TS-derived CV, reweighted FES projected along funnel axis distance and ligand solvation and multiple metastable states in the transition pathways for OAMe-G1, OAMe-G4 and Trypsin-Benzamidine complexes respectively. For(D-F) Free energy convergence plots in different time intervals for three different complexes. Free energy at different time intervals is colored with different colors.

Next we have investigated the mechanism of ligand binding in Trypsin-Benzamidine complex are the results (time series, FES, and configurations of different states) are presented in Figure 3C. For this complex also, bound, unbound and an intermediate – three metastable states emerges in the reweighted FES shown in Figure 3C. A barrier of ∼19 KJ/mol is present going from intermediate to unbound conformation. Time series plot of dist.z shows benzamidine ligand goes from bound conformation (dist(z) ∼ 0.22-0.38 nm) to unbound conformation (dist(z) ∼2.35-2.6 nm) quite frequently within 200 ns, demonstrating the efficacy of TS-derived CV. The different metastable conformations along the transition pathway are shown in Figure 3C. Free energy convergence plot shows FE along dist.z converges after just 60 ns. For the same system, using more powerful OPES method, it requires almost three times more sampling time to converge the whole FE landscape.(26)This clearly demonstrates the strength of TS-derived CV to explore the complex FES rapidly and attaining convergence of free energy on relatively short timescale. For trypsin-benzamidine complex, we have shown the water density around the ligand in different metastable states in Figure 3C. The change is water density around the ligand, benzamidine is well depicted, highlighting the crucial role of ligand solvation along the transition pathway. We also run WT-MetaD with just funnel axis distance for every ligand complexes. It is observed that the number of transitions between bound and unbound conformation is reduced compared to those observed when biasing with TS-derived CV. For a fair comparison, we have used same height and biasfactor for both the two cases. The trajectories biasing with only funnel axis distance is shown in Figure S7A. Moreover, we also check the convergence of free energy when biasing with funnel axis distance. Although for solvent exposed ligand complexes, free energy converges with funnel axis distance but it takes much longer time to converge compared with TS-derived CV. The free energy convergence plot biasing with only funnel axis distance is shown in Figure S7B. The difference is more pronounced in case of Trypsin-Benzamidine, where WT-MetaD along funnel axis distance takes almost 200 ns to converge in contrary of just 65 ns in case of TS-derived CV indicating almost five times acceleration in free energy convergence.

### (B) Case of semi-buried and buried active site

Next we move towards more challenging systems which have buried active site to investigate the binding mechanism. We have chosen three systems: (a) L99A T4 Lysozyme-benzene complex (buried active site), (b) HSP90 ligand complex (semi-buried active site) and (C) Soluble sodium epoxide hydrolase ligand complex (buried active site). Similar to solvent exposed case, we have first chosen three OPs to describe the binding-unbinding transitions here. Interestingly, analysis of the OPs near TS region from explainable AI model reveals that active site solvation has almost zero contribution for each system studied here. It is somewhat physically intuitive, as in buried ligand-bound complexes active site solvation does not change noticeably when binding or unbinding event occurs, therefore contributing very minimally to ligand binding transition dynamics. We have evaluated the contribution of different OPs near the TS using the methodology described before for L99A T4 Lysozyme benzene complex and shown in Figure 4A. The coefficient of ligand solvation is quite high (0.98) compared to that of COM distance (0.22). This indicates that ligand solvation is majorly driving the binding/unbinding transition in buried active site complexes, consistent with our previous observations for solvent exposed ligand complexes. We also evaluated the relevance of different OPs near TS for other two systems and given in Figure S8. Interestingly, for both complexes we found ligand solvation coefficient is much higher than the distance similar to Lysozyme benzene complex. Although the derived coefficients of OPs for each complex are slightly different, however relative order of coefficients remains same. Now, to use one transferable CV for all the buried/semi-buried systems we directly use CV constructed from the T4L lysozyme benzene complex to other two complexes. We have shown the bound conformations for each system studied here in Figure 4(B).

**Figure 4:**
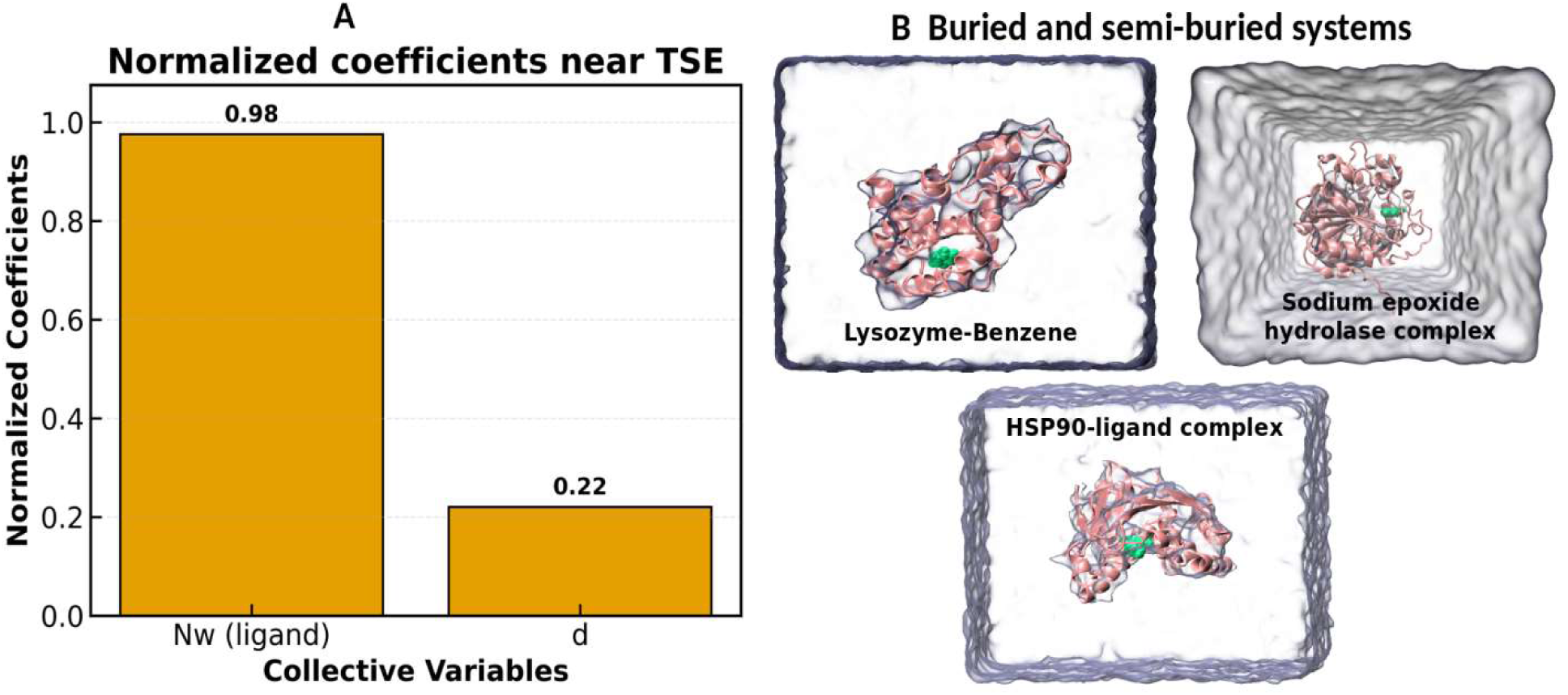
(A) Normalized coefficients of OPs near TSE for buried active site protein-ligand complexes. (B) Bound conformations of three buried systems used in this work.

We run WT-MetaD for three complexes with the TS-derived CV constructed from interpretable ML model. We first investigate the mechanism of ligand binding T4L lysozyme complex which has deeply buried active site. The reweighted FES shown in Figure 5A demonstrated the existence of multiple metastable states along the transition pathway and representative structures of each minima is shown. Time series plot along distance for this complex suggests multiple back and forth transitions between bound and unbound conformations happens within 250 ns. Interestingly for this complex, a bound like conformation (distance ∼0.55 nm) is present in the FES where ligand solvation is slightly higher than that of bound conformations (Figure 5B). Going from this bound-like conformation to bound conformation, a substantial barrier of nearly 24 KJ/mol is present. This suggests along this binding pathway 4-5 water molecules expelling from the ligand is necessary to reach the final bound conformation. Convergence plot for this complex shown in Figure 5D depicts that FE reaches convergence within just 36 ns.

**Figure 5:**
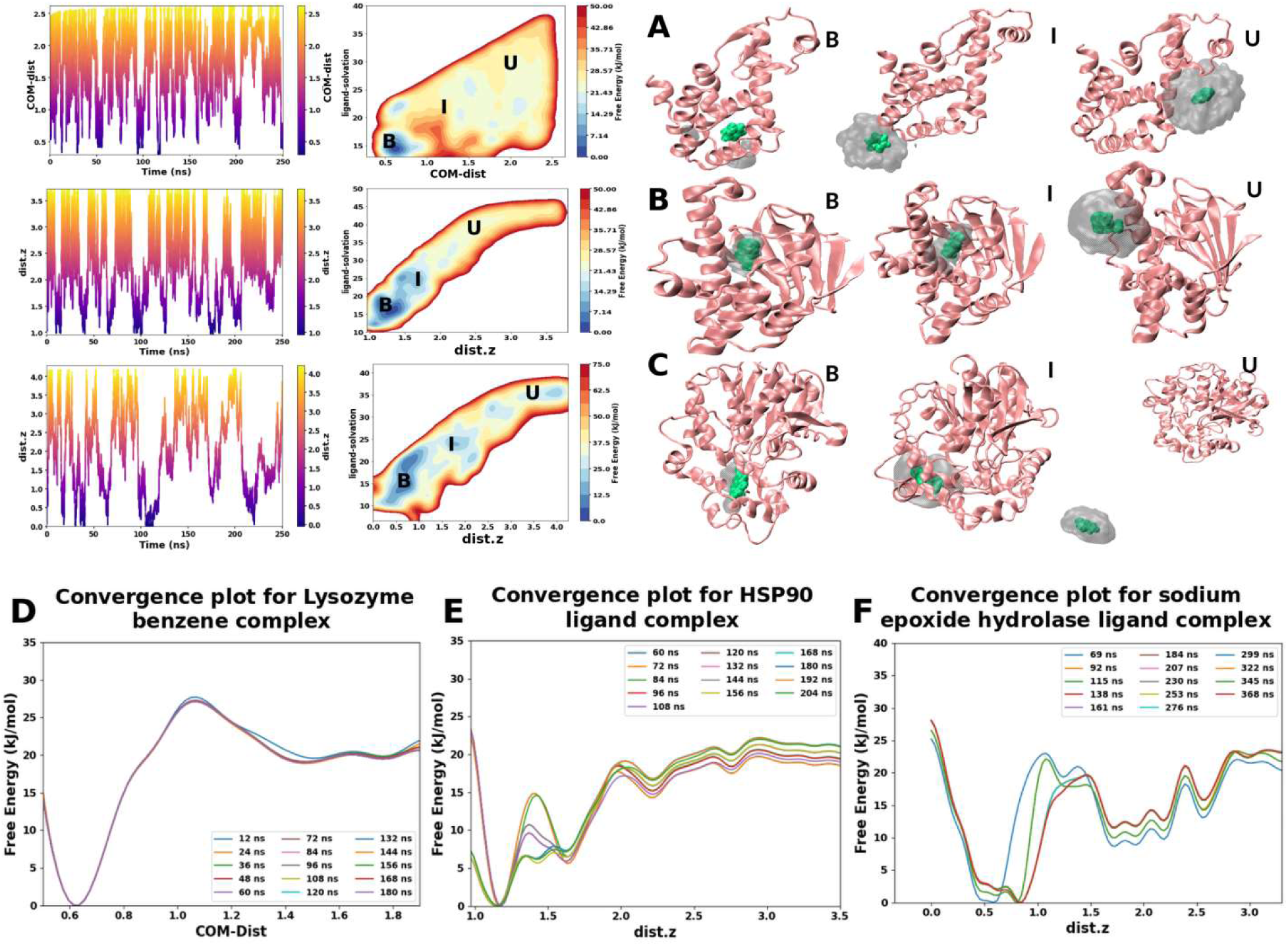
(A-C) Trajectories of WT-MetaD biased with TS-derived CV, reweighted FES projected along funnel axis distance and ligand solvation and multiple metastable states in the transition pathways for lysozyme-benzene complex, HSP90 ligand complex and sodium epoxide hydrolase ligand complex respectively. (D-F) Free energy convergence plots in different time intervals for three different complexes. Free energy at different time intervals is colored with different colors.

Next we tested the efficacy of TS-derived in CV in semi-buried HSP90 ligand complex. We have shown the reweighted FES in Figure 5B. A FE barrier of ∼ 18 KJ/mol is present between bound and unbound conformation. Representative structures of each conformation are shown in Figure 5B. Time series plot of distance shown that large number of re-crossing events of binding happens within very short time. The FE convergence plot for HSP90 ligand complex is shown in Figure 5E where FE converges within 120 ns biasing with TS-derived CV.

Finally, transferable TS-derived CV is tested in more challenging system, sodium epoxide hydrolase ligand complex which has extremely deeply buried active site. Reweighted FES presented in Figure 5C looks much more rugged landscape as multiple metastable basins present other than just bound and unbound conformations. Interestingly for this complex also, our simulation discovered an additional bound like conformation which has dist.z value similar to that of bound conformation, but a slight change in hydration water of ligand (∼4-5 water molecules) is present. Time series plot suggests that multiple back and forth transitions between dist.z (∼0.05-0.5 nm) to dist.z (∼3.5-4.1 nm) happens even for this complex. The convergence of FE analysis depicts that for this complex, FE reaches a plateau after 200 ns which is slightly higher than all the other complexes studied here (Figure 5F). It should be noted that buried active site protein complexes, ligand binding-unbinding process is more challenging than that of solvent exposed systems as the former requires movement of the ligand to the surface of the protein and then tunnel through the protein to reach the final active site. In a similar way, as in case of solvent exposed systems we have conducted WT-MetaD with only funnel axis distance/COM distance for these buried active site complexes. Interestingly, for these complexes binding-unbinding transitions with distance based CVs is much lower than that of TS-derived CV. Trajectories biasing with only distance based CV for each buried active site complexes are given in Figure S9. The free energy convergence plot biasing with distance are given in Figure S10, where it is demonstrated that convergence took much longer time than that of TS-derived CV. Moreover, for sodium epoxide hydrolase ligand complex we do not even get the free energy convergence using funnel axis distance as a biasing CV. This highlights the need of incorporating ligand solvation explicitly in deriving optimal CV to investigate ligand recognition in buried active site complexes.

### (C) One CV to enhance the sampling of ligand binding transition

In this section, we aim to propose a transferable CV to investigate binding-unbinding transitions in different class of protein ligand complexes. It is already shown that for both solvent exposed and buried ligand bound complexes, ligand solvation emerges to be the dominant coordinate in driving transitions. However, active site solvation, comes as a crucial coordinate only in case of solvent-exposed systems, for buried complexes the contribution of active site solvation at the TSE is negligible. Therefore, to propose a transferable CV for every ligand complexes we use TS-derived CV from buried system to test its efficacy in solvent-exposed complexes. Interestingly, when buried active site TS-derived CV is used in solvent-exposed systems, multiple back and forth transitions is observed between the metastable states as similar to that of with TS-derived CV from solvent exposed cases. This demonstrates that a CV where ligand solvation plays the most dominant role, can trigger transitions for ligand complexes irrespective of their nature of active site. We have shown the reweighted FES along funnel axis distance and active site solvation in Figure 6(A). Interestingly, for OAMe-G1, OAMe-G2 and OAMe-G4 systems, there are two alternate pathways of ligand binding as depicted in Figure 6A. One pathway is dry pathway, where active site solvation does not change much during unbinding and another pathway is wet pathway where both funnel axis distance and active site solvation sharply changes during unbinding. Intriguingly, dry pathways for each OAMe-guest systems are high energy pathway and wet pathway follows the minimum free energy pathway (dominant one). This result is also consistent with the previous studies, but these studies included active site solvation for biasing to construct the converged FES. However, we get the exact converged FES along funnel axis distance and active site solvation without even taking into account the active site solvation degree of freedom to explore the FES. For hydrophobic cavity-fullerene system and trypsin benzamidine, we do not find any signature of dry pathway in the FES of ligand binding (Figure 6A). We have calculated binding free energy for each complex by funnel correction and shown the correlation plot of estimated binding free energy from our calculation and the experiments in Figure 6B. Encouragingly we got quite good correlation between the estimated binding free energy and experiments. For all the ligand complexes binding free energies are within 1Kcal/mol range relative to the experimental values. In case of host-guest systems and HSP90 ligand complex, computed binding free energies are within 0.30 Kcal/mol range relative to the reference values. For L99A T4 lysozyme complex obtained binding free energy is 0.45 Kcal/mol lower than that of experimental values. For trypsin benzamidine complex and sodium epoxide hydrolase ligand complex, the estimated binding free energies differs 0.65 and 0.70 Kcal/mol respectively from the experiments. For trypsin benzamidine complex, Rizzi et al. calculated binding free energy with taking into account different hydration site collective variable in the protein and got exact match with the experiments.

**Figure 6:**
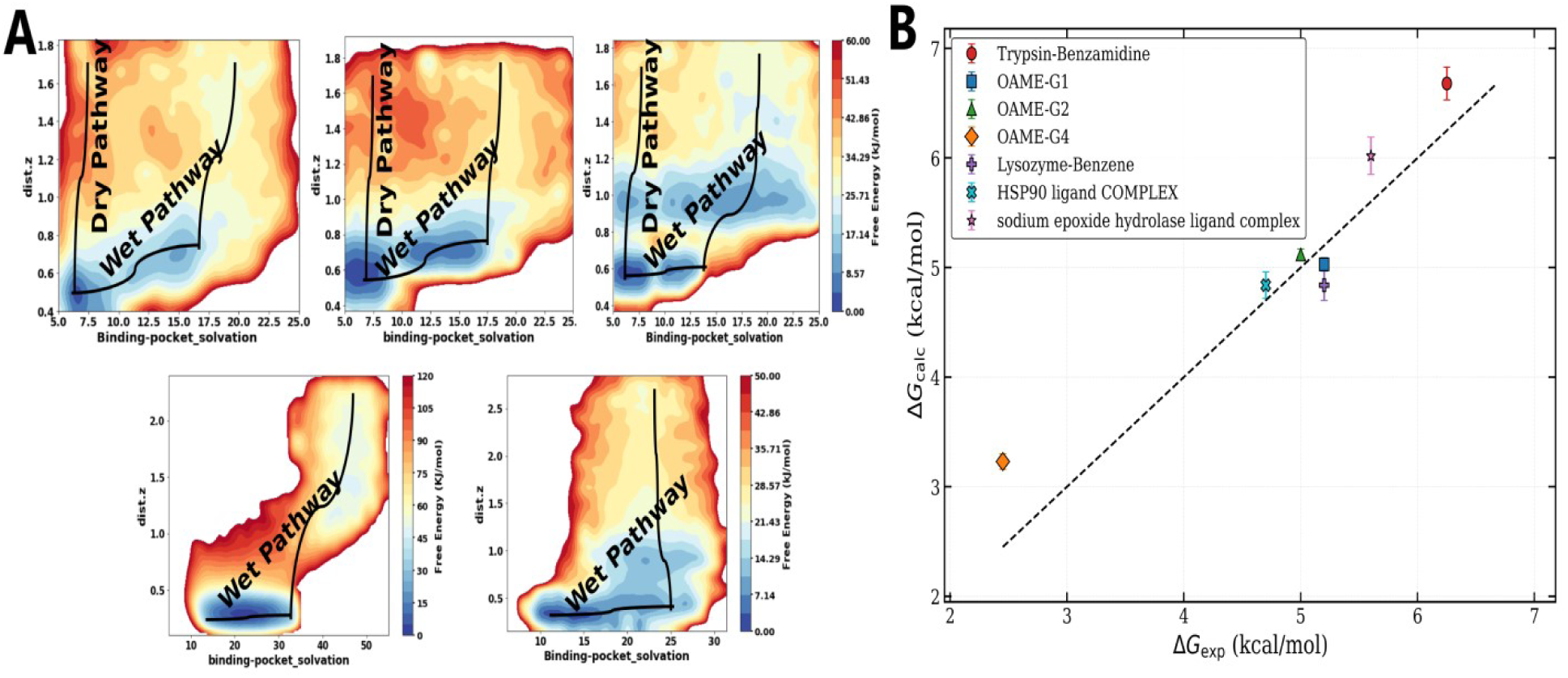
(A) Reweighted FES along funnel axis distance and binding site solvation for solvent exposed active site complexes biasing with TS-derived CV for buried active site systems. (B) Correlation plot of estimated binding free energy and experimental binding free energy for all the protein-ligand complexes studied in this work.

### (D) Drug Residence time calculation using transferable TS-derived CV and infrequent metadyanmics

Accurately estimating the drug residence times hold great promise in drug discovery pipeline. The final transferable TS-derived CV was used to estimate the residence times of different ligands with different hosts studied in this work. We have used Transferable TS-derived CV to run multiple independent iMetaD runs. From these iMetaD runs, acceleration factors are calculated and residence times are estimated. The whole workflow of residence time calculation is given in Figure 7(A). The poison fit of the reweighted transitions for OAMe-G2, HSP90 ligand complex and sodium epoxide hydrolase ligand complex are given in Figure 7(B). The p-values are estimates using Kolmogorov-Smirnov test (KS-test). P-values for all the complexes are greater than 0.05 justifying the reliability of the estimated residence times. P-values for HSP90 ligand complex is relatively lower as depicts in Figure 7(B), suggesting multiple transition pathways associated for this complex. The poison fit of the other complexes are given in Figure S11. In Figure 7C, the estimated residence times and reference residence times (experimental or computationally computed) are given which agrees quite well with the reference value. For OAME-complexes authors from Ref. estimated residence time using GAMBESS and the estimated times from our methods exactly matches. For L99A T4 lysozyme-benzene complex, we get different residence time depending upon the transition path which is consistent with the previous studies. However, we get the fastest timescale of ∼9.6 ms which closely matches with the experimental timescale of 1.52 ms. For hydrophobic cavity fullerene system we get the residence time of about ∼5000 s which closely matches with the experimental value of ∼4000 s. For trypsin-Benzamidine complex, we get residence time of about 7.8 X 10^−3^s which aligns within reasonable range of 1.6 X 10^−3^ s residence time of experiments. For sodium epoxide hydrolase ligand complex and HSP90 ligand complex studied here, we get residence time of 1.04 X 10^−2^s and 7 s, respectively. We do not find experimental or computational residence time for these two complexes to directly compare with our method. the Further experimental and computational studies are needed to carefully estimate the kinetics for these systems. Nonetheless the proposed protocol accurately estimates the kinetics for six ligand bound complexes studied here. This highlights that this protocol can be extended to real drug discovery pipeline and to do initial virtual screening for different new target proteins to rule out inefficient drugs/inhibitors whose residence time is extremely low.

**Figure 7:**
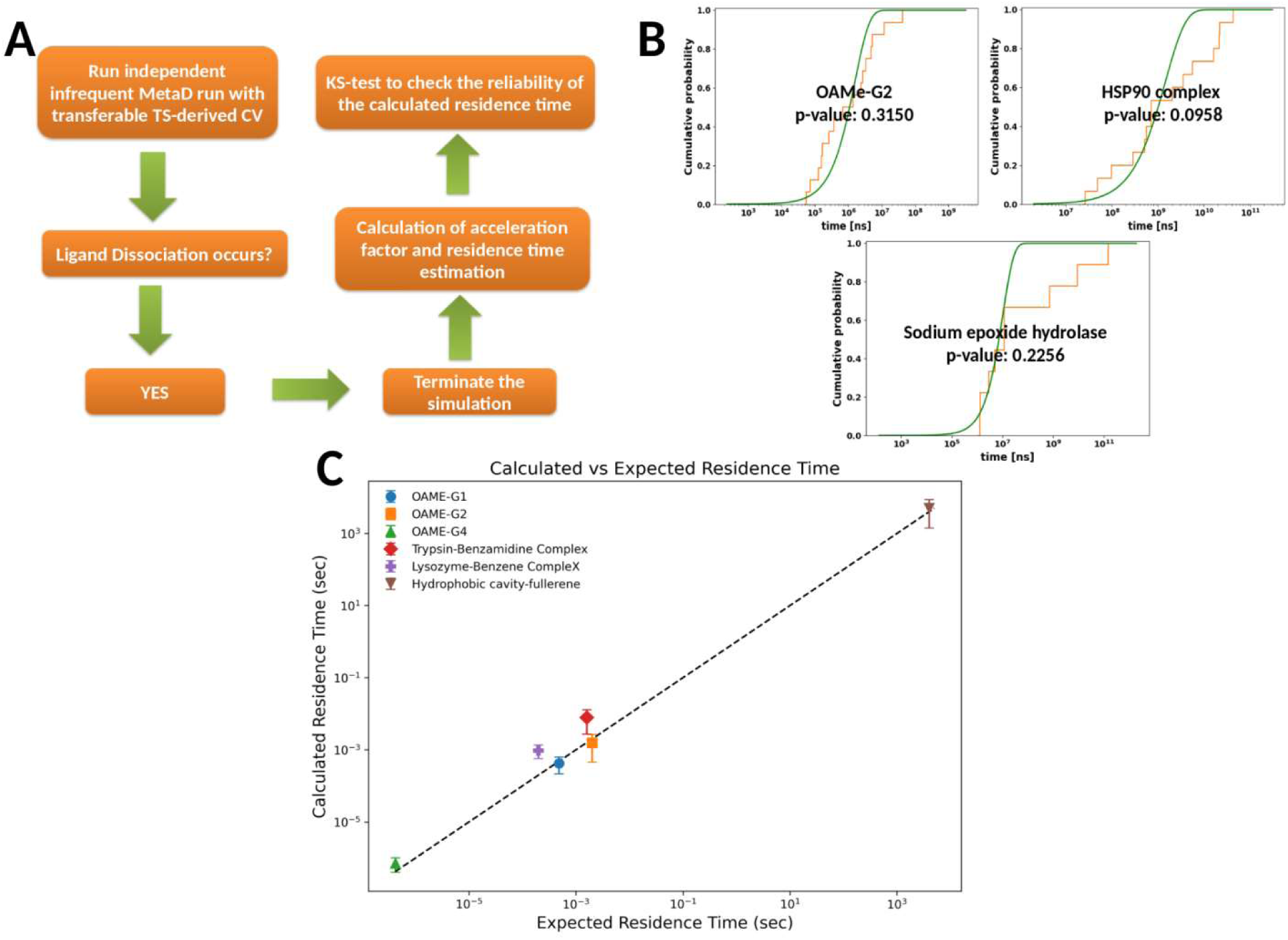
(A) Protocol to estimate residence times of ligands using TS-derived CV with iMetaD, (B) Cumulative distribution curve for three ligand complexes and associated p-values and (C) Correlation plot of estimated ligand residence time from TS-derived CV with reference values.

## Conclusion

Investigating the mechanism of drug-receptor binding holds immense significance inmodern drug discovery. Enhanced sampling methods are widely used to accelerate the FES exploration of complex drug binding mechanism, their efficiency heavily relies on the design of optimal CVs. Designing an appropriate CV typically requires extensive system-specific knowledge to accurately capture the slowest degrees of freedom. In this study, we propose an explainable AI driven framework designed to derive optimized CV from TSE for diverse ligand bound complexes. The TS-derived CV is shown to be extremely effective in accelerating ligand binding transition and achieves faster free energy convergence in case of both solvent exposed and buried active site systems. Intriguingly, ligand solvation is found to be the most crucial coordinate in driving binding transition while analyzing the TSE in both scenarios. Most importantly, we propose a transferable TS-derived CV which can be used across diverse ligand bound complexes to elucidate small molecule recognition mechanism. This transferable framework drastically reduces the need of incorporating system specific CV and eliminates the requirement of new training data to design ML-derived CV when investigating entirely new systems. Moreover, we also integrate this transferable TS-derived CV with iMetaD to estimate the ligand residence time across all the studied complexes and interestingly estimated residence time matches closely with that of reference residence times.

Overall, this framework provides a less computationally demanding yet accurate approach for estimating binding free energy and ligand residence time, opens up new opportunities to screen large library of ligands in a virual screening workflow and direct integration into drug design workflow. This framework can also be integrated with automated sophisticated clustering methods to decode multiple ligand binding transition pathways and different metastable states. For kinetics estimation, we use iMetaD method, however this can be replaced by any other new variant of infrequent metadynamics or OPES-based methods. Despite the strength of this current protocol in accelerating transition and getting reliable thermodynamics and kinetics associated with the system, challenges remain that need addressed in future. First of all, it should be noted that this framework can be only applicable for the protein-ligand complexes which do not undergo large scale conformational transition as explicit conformational coordinate is not incorporated in the current framework. Second, prior knowledge of the target binding pocket remains a prerequisite for investigating the binding mechanism. In the absence of the knowledge of active site, designing a complete blind exploration protocol without any a-priori knowledge of known active site has to be developed. Thirdly, in case of flexible target bio-molecules (like protein-peptide, nucleic acid-peptide and protein-protein recognitions) where structural plasticity of the constituents will pose additional difficulties to decipher the mechanism of ligand binding. In future, we will work on these directions.

## Data Availability

The input files for performing all the simulations in this work and code to find out the contributions of different order parameters near the transition state are given in the GitHub repository: https://github.com/saikat-ai/ligand_binding_role_of_ligand_solvation.

## ACKNOWLEDGMENTS

We would like to thank ANRF (Grant no: ANRF/ARG/2025/003888/CS) and Indian Association for the Cultivation of Science (IACS) for computational facilities. S.A thanks IACS for the fellowship. S.A thanks Mr. Priyabrata Dey and Rohon Mitra for insightful discussions.

## Author Contributions

B. J. designed research; S.D. performed research, S.D. analyzed data; S.D. and B.J. jointly wrote the paper.

## Supporting Information

Details of infrequent metadynamics, funnel metadynamics, chosen biasfactor and Gaussian height for performing metadynamics simulation, Trajectories biasing with distance in different ligand complexe, FE convergence plot biasing with distance and cumulative distribution plots of residence times for different ligand complexes.

---

Note S1: Details of infrequent metadynamics and funnel metadynamics:

## Infrequent metadynamics

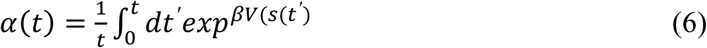

In the above equation, V (s, t′) is the metadynamics time-dependent bias, 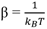 is the inverse temperature multiplied by the Boltzmann constant k_B_.Here, α(t) is the accelerated time. A single infrequent MetaD simulation is not sufficient to calculate kinetics for any transition, generally multiple infrequent metaD simulation runs are conducted to estimate kinetics. The simulations were truncated as soon as the end state is reached. In this study, starting state is bound one and end state is unbound conformation. The series of ligand residence times obtained from multiple independent infrequent metaD runs should follow poisson distribution. The quality of estimated kinetics is checke via KS-test. P-value greater than 0.05 in the KS test indicates that the residence time does not significantly deviate from the Poisson process model expected for rare events. This suggests that the estimated accelerated timescales can be trusted.

## Funnel metadynamics

In a standard MetaD simulation once the ligand leaves the binding site, it enters into bulk solvent region where it can diffuse freely. As the search space in the solvent is huge, the ligand spends a large amount of time exploring the non-relevant region rather than again coming back to the actual binding site. This poses challenge to converge the MetaD simulation and accurate estimation of the binding free energy. Therefore to mitigate this issue, funnel metadynamics applies a cylindrical restrain potential which restricts ligand’s motion into a narrow volume which includes binding site and a path connecting to binding site and bulk solvent. Interested readers can read detailed discussion on funnel MetaD in the following references. For the funnel restraint potential, free energy difference between bound and unbound state needs an entropic correction and can be calculated from free energy profiles using the following expression:

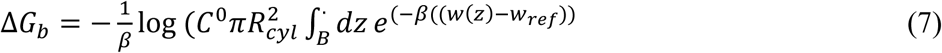

In this equation and 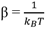 kB is the boltzman constant, R_cyl_ is the radius of the cylindrical region of the funnel. W(z) is the free energy along funnel axis, W_ref_ is the reference free energy value in the unbound state 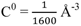 is the standard concentration. The integral is performed over the bound (B) conformation region.

---

**Table S1:** Chosen sigma of TS-derived CV for different scenarios:

| Different ligand binding systems | Sigma for TS-derived CV |
| --- | --- |
| Solvent exposed ligand bound complex | 0.12 |
| Semi-buried/ Buried bound complex | 0.04 |

**Table S2:** Chosen biasfactor for each system to run MetaD with TS-derived CV.

| Systems | Biasfactor | Height |
| --- | --- | --- |
| Hydrophobic cavity fullerene | 30 | 1.5 |
| OAMe-G1 | 20 | 1.4 |
| OAMe-G2 | 20 | 1.4 |
| OAMe-G4 | 20 | 1.4 |
| Trypsin-Benzamidine | 30 | 1.5 |
| HSP90 complex | 30 | 1.5 |
| Lysozyme Benzene complex | 20 | 1.5 |
| Sodium epoxydase complex | 30 | 1.5 |

**Figure S1:**
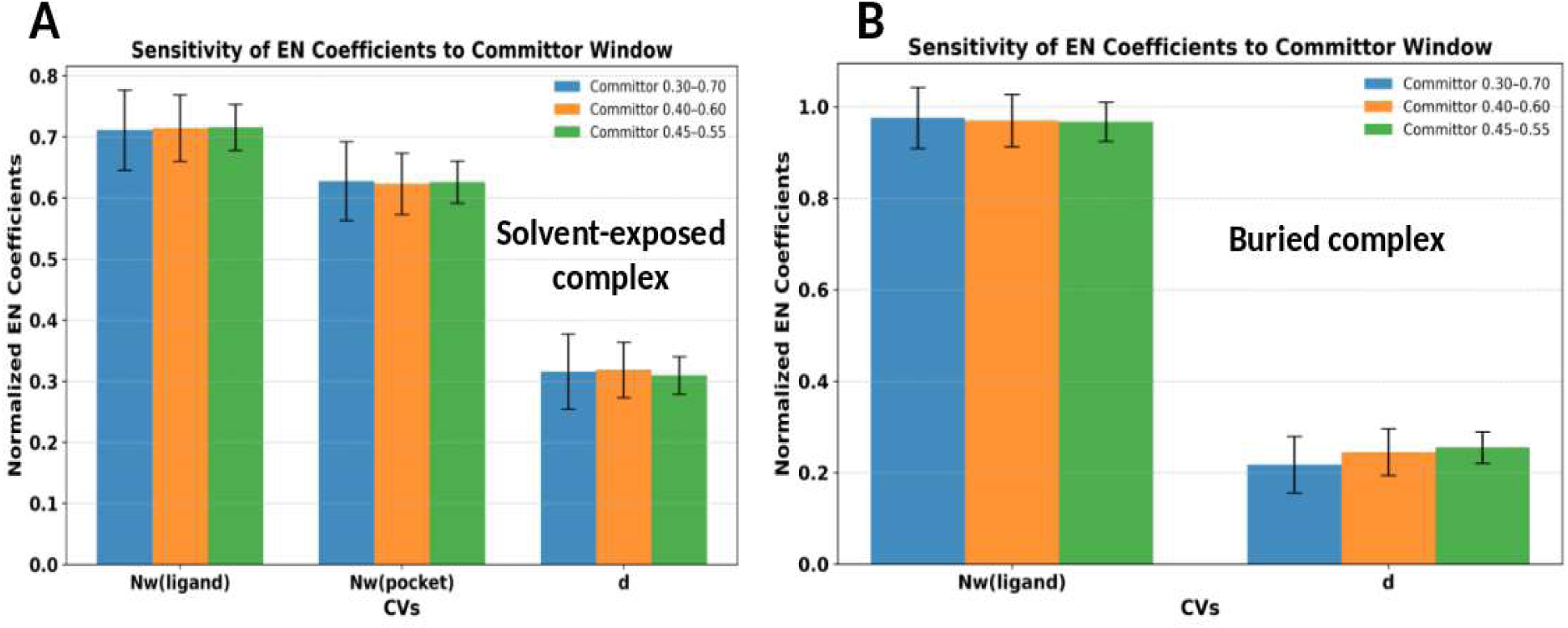
Normalized coefficients of different OPs in two different binding scenarios in selected committor ranges:

**Figure S2:**
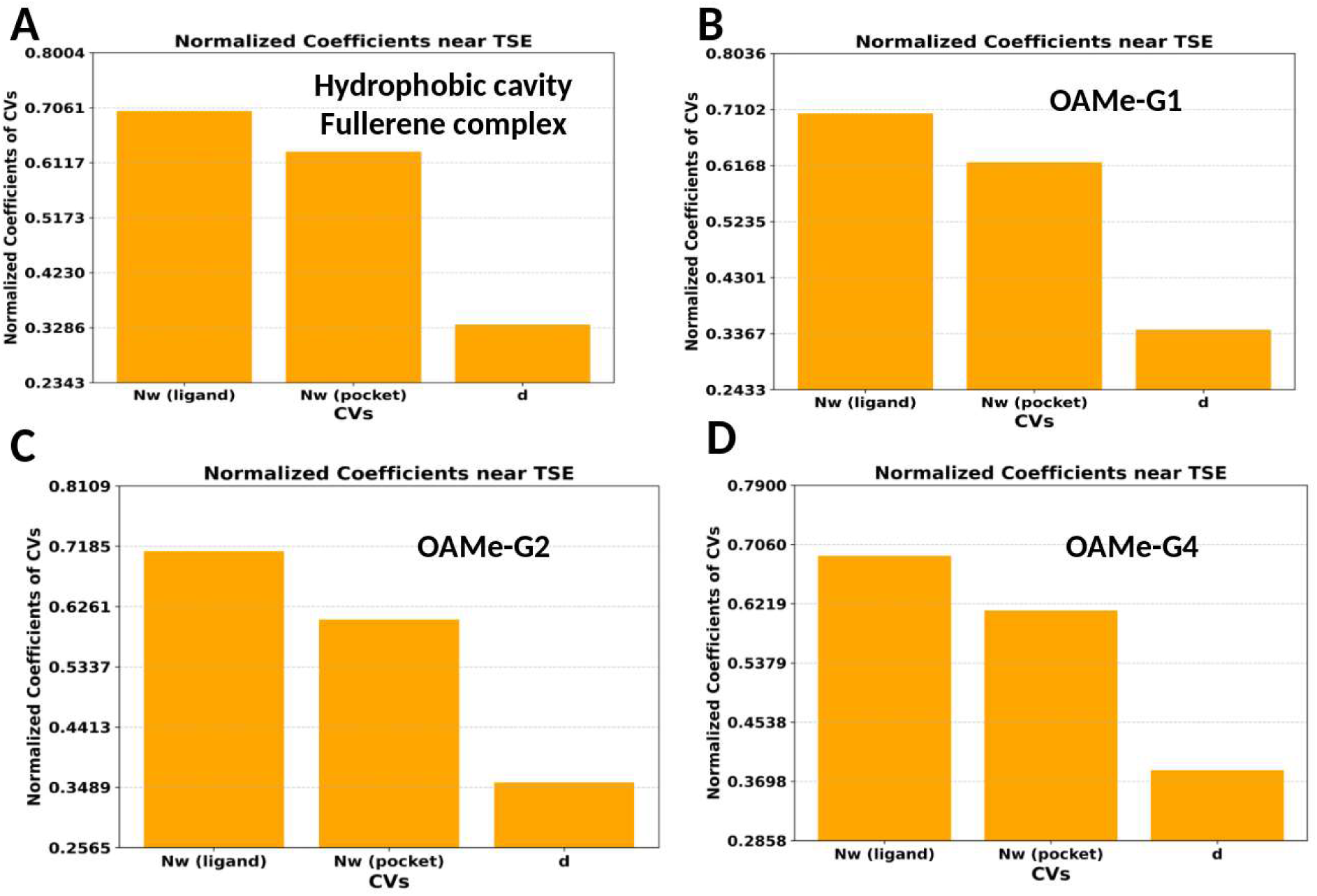
Normalized coefficients of different OPs for solvent exposed systems near TSE:

**Figure S2:**
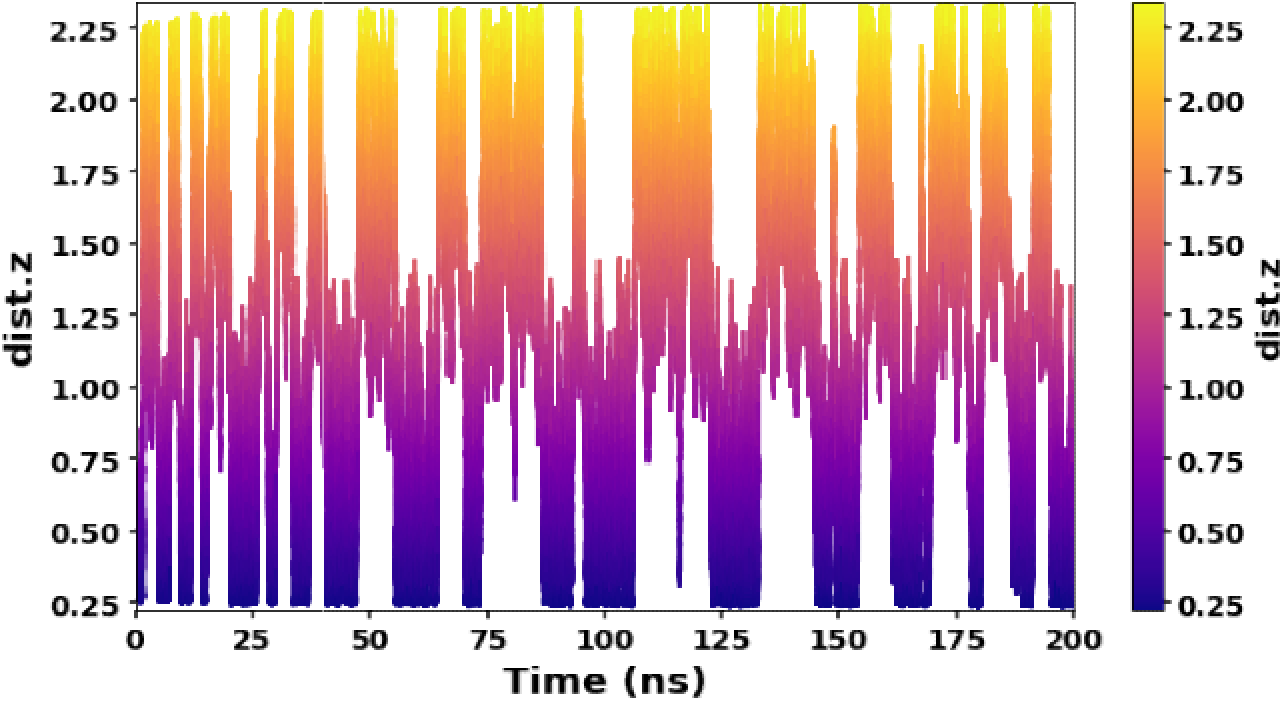
Time series plot of dist.z during 200 ns MetaD simulation with TS-d hydrophobic fullerene system rived CV for

**Figure S3:**
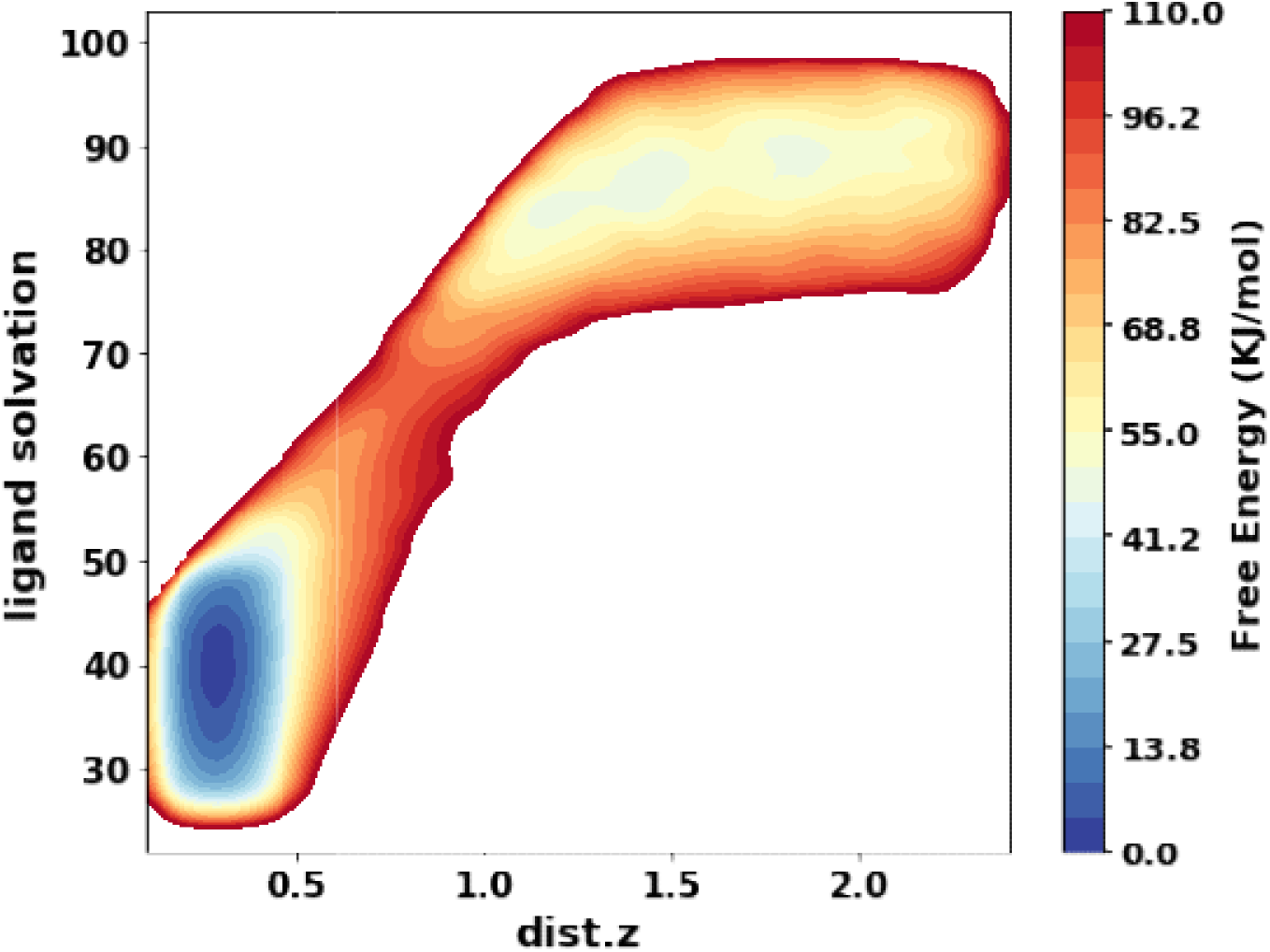
Reweighted FES plot of dist.z and ligand solvation for hydrophobic fullerene system

**Figure S4:**
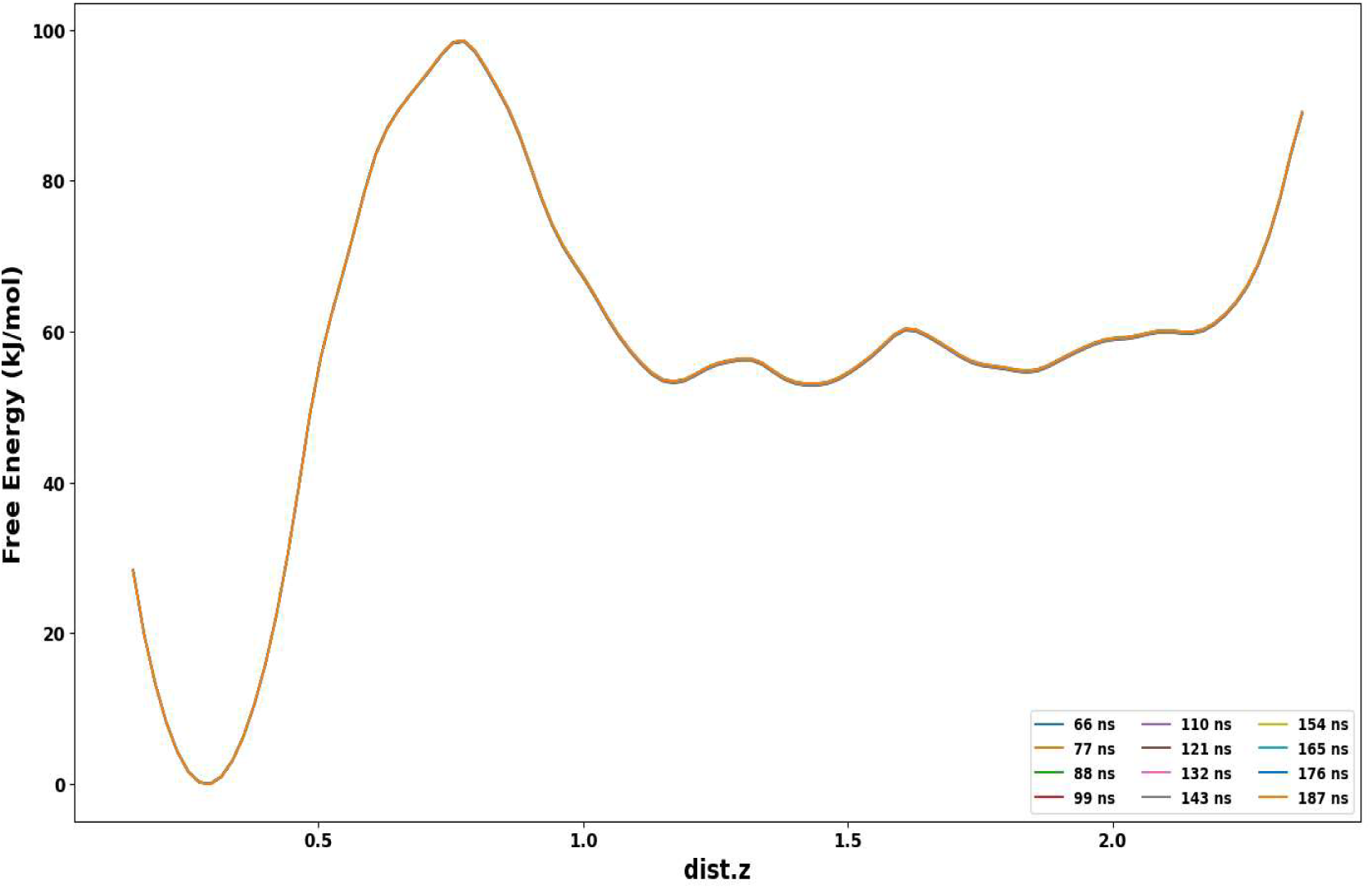
Free energy convergence plot of dist.z in hydrophobic cavity system

**Figure S5:**
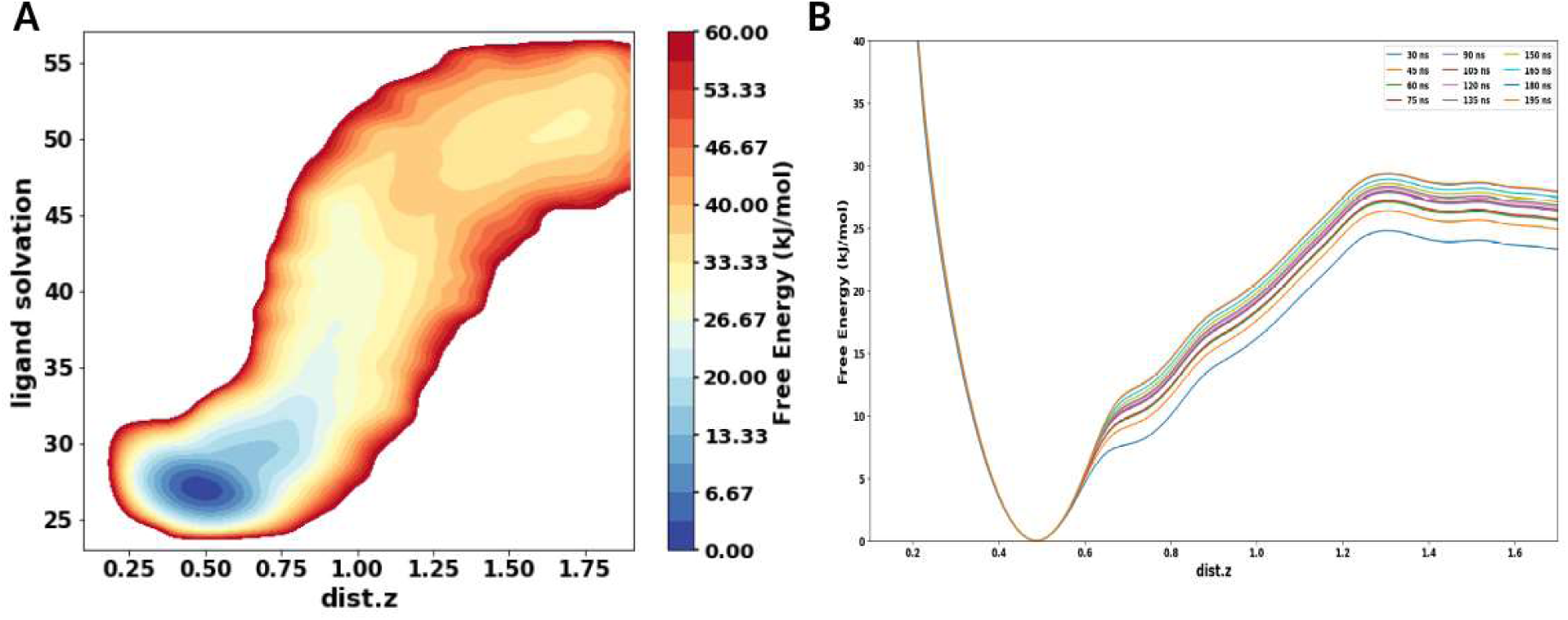
(A) Reweighted FES along dist.z and ligand solvation biasing with funnel axis distance for OAMe-G1 and (B) Free energy convergence plot for OAMe-G1.

**Figure S6:**
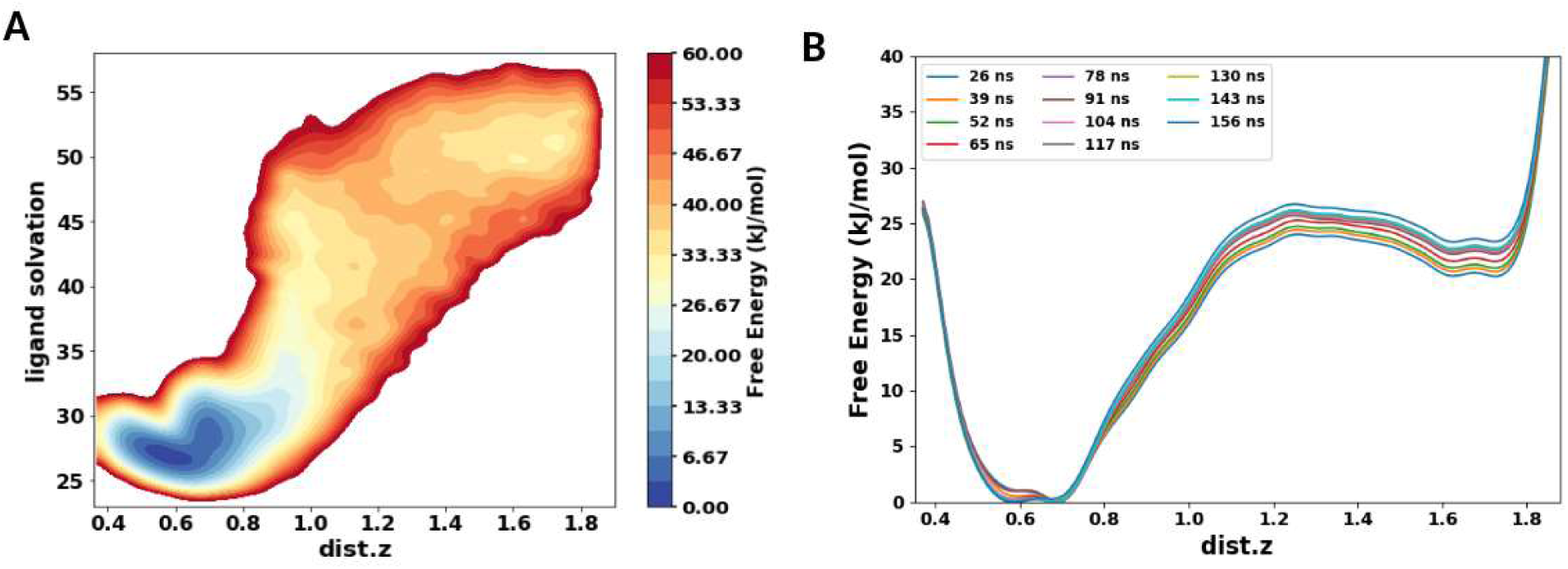
OAMe-G2 complex (A) reweighted FES along dist.z and ligand-solvation and (B) FE convergence plot

**Figure S7:**
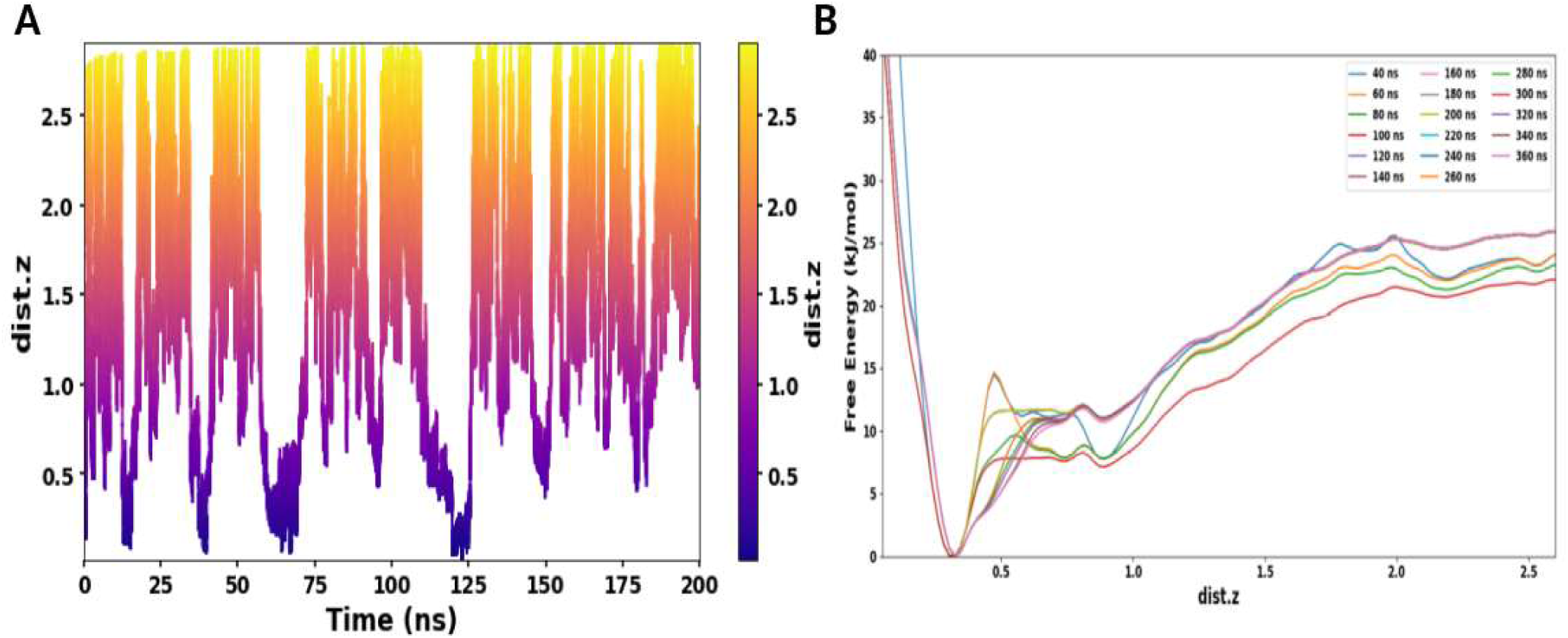
(A) Time series plot of dist.z biasing with funnel axis distance in Trypsin-Benzamidine system and (B) Free energy convergence plot biasing with funnel axis distance

**Figure S8:**
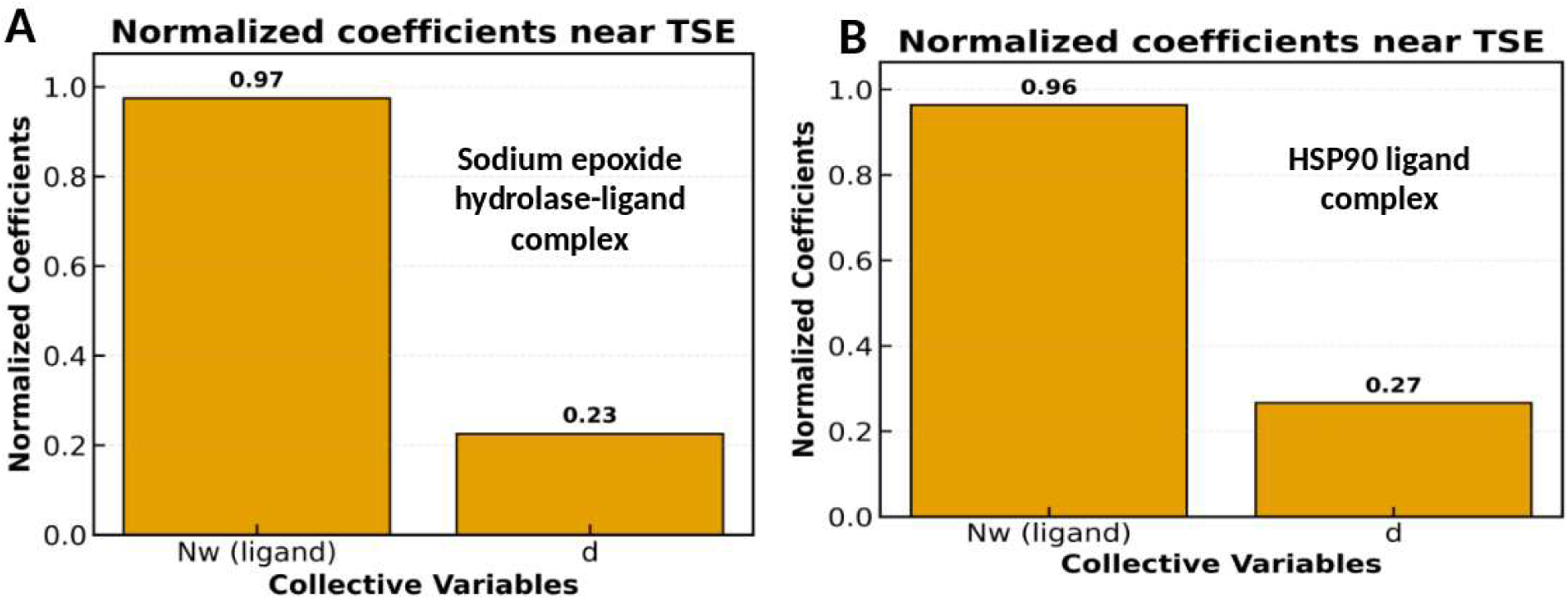
Normalized coefficient of different OPs near TSE for (A) sodium epoxide hydrolase ligand complex and (B) HSP90 ligand complex

**Figure S9:**
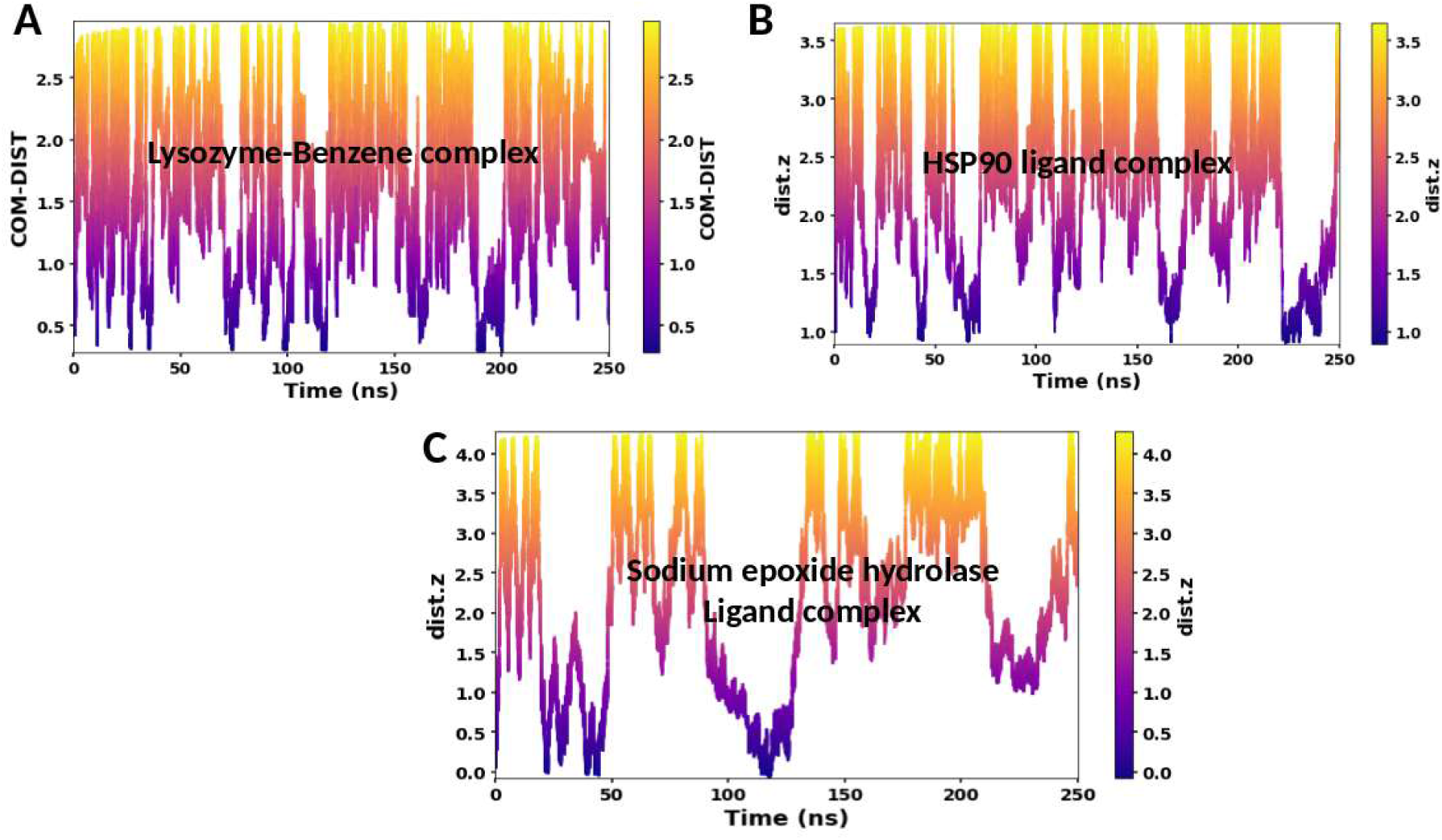
Trajectories biasing with distance in case of active site buried complexes:

**Figure S10:**
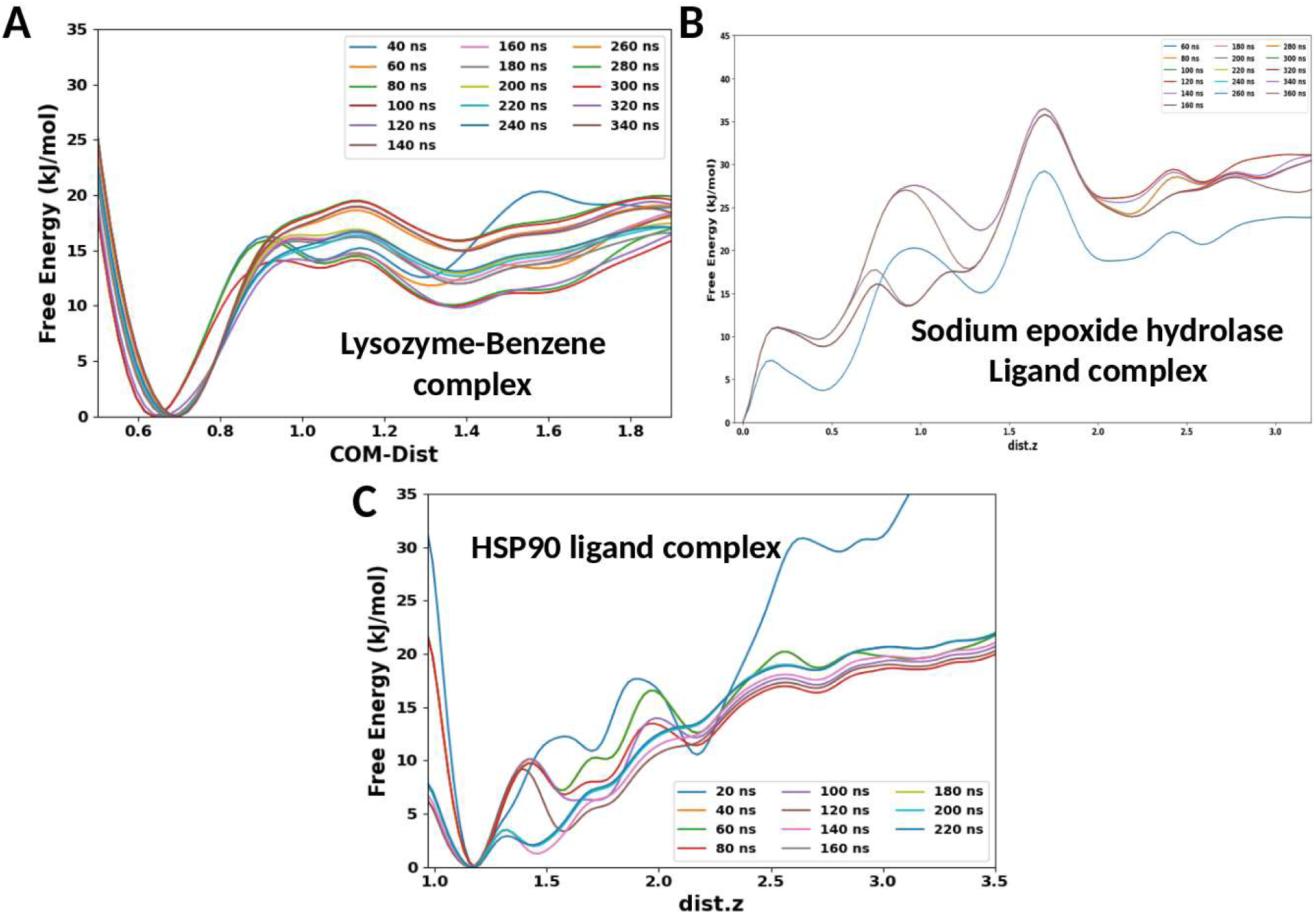
FE convergence plot for three buried active site complexes biasing with distance

**Figure S11:**
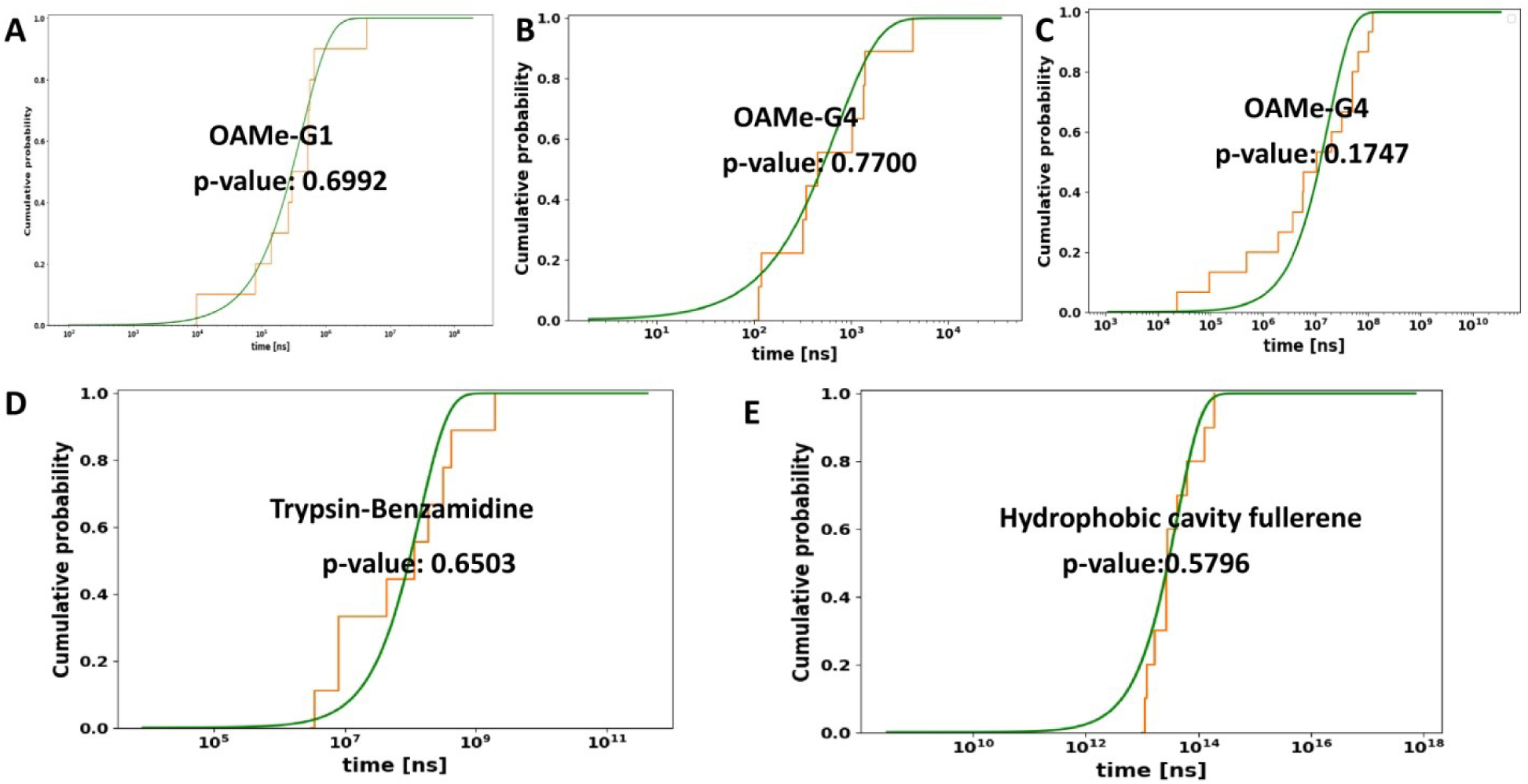
Cumilative probability distribution plot of residence times for other complexes

